# Novel insights into conserved biomineralization mechanisms revealed from a cold-water scleractinian coral skeletal proteome

**DOI:** 10.64898/2026.03.24.713908

**Authors:** Jeana L. Drake, Tali Mass, Liti Haramaty, Erik E. Cordes, Santiago Herrera, Paul G. Falkowski, Oded Livnah, Fiorella Prada

## Abstract

Stony corals exhibit striking morphological plasticity across diverse environments and trophic strategies, raising fundamental questions about the conservation of their biomineralization machinery. Here, we characterize the skeletal organic matrix proteome of the cold-water, asymbiotic coral *Desmophyllum pertusum* and compare it with skeletal proteomes from a facultatively photosymbiotic temperate coral and an obligately photosymbiotic subtropical coral. Despite pronounced differences in habitat, symbiotic status, and skeletal micro-density, we observe convergence on a conserved “biomineralization toolkit”. Comparative proteomics, genomics, and structural predictions reveal that this toolkit integrates diverse acidic matrix proteins, carbonic anhydrases, adhesion and structural proteins, and signaling components with multiple cellular export pathways. Together, these findings redefine coral biomineralization as a dynamic, coordinated network of cellular pathways rather than a static assemblage of matrix components. The symbiont-free model *D. pertusum* provides a mechanistic framework for dissecting coral calcification across environments and for assessing the resilience of this conserved machinery.

## Introduction

Stony (scleractinian) corals are cnidarians that produce external calcium carbonate skeletons, usually of the polymorph aragonite ^1^. In colonial species, these skeletons form a complex reef framework that provides habitat for ∼25% of described marine fish and invertebrate species globally ^2^. Remarkably, such reef-building capacity spans a wide range of environmental conditions, from warm, shallow, sunlit waters to cold, dark, food-limited deep seas ^3^.

Despite profound differences in temperature, light availability, trophic strategy, and symbiotic status between shallow- and cold-water corals, multiple lines of evidence indicate conserved mechanisms of skeletal formation. Genetic comparisons ^4^, similarities in tissue-skeleton interface ^5,6^, and shared macro- and micro skeletal structures (e.g.,^7,8^ suggest that a common biomineralization machinery underlies coral calcification across habitats. Yet, how this machinery operates at the cellular and molecular levels, and how it remains effective across such disparate environments, remains largely unresolved.

Across taxa, biomineralization generally occurs within a defined physical space whose chemistry is tightly regulated by biomolecules. In stony corals, adjacent cells (e.g., calicoblastic cells) produce biomolecules ^9^ that sequester dissolved inorganic ions into a "privileged space”, commonly identified as the extracellular calcifying medium (ECM) ^10^. The ECM contains seawater-derived ions ^11^, supplied in part via paracellular transport across intercellular junctions ^5,12–14^. The ECM chemistry is also actively modified through (a) transmembrane ion transporters such as Ca-ATPase pumps and bicarbonate anion transporters ^15,16^, (b) macropinocytosis of ECM into calicoblastic cells where it is likely modified through exchanges with the cytosol ^13^, and (c) intracellularly-derived vesicles that accumulate cytosolic and organellar materials ^5,14,15,17^.

In addition to regulating the inorganic chemistry, the calicoblastic cells secrete a skeletal organic matrix (SOM) composed of a mixture of polysaccharides, lipids, and proteins ^18^. These macromolecules can drive the nucleation, growth, and inhibition of the mineral phase in particular directions and influence biomineral mechanical properties ^19^. Indeed, the organic component, although minor by weight, enhances fracture resistance by orders of magnitude relative to the pure mineral, as demonstrated across diverse biomineralizing taxa ^20^. Among SOM molecules, proteins are the most extensively studied across biomineralization taxa ^21,22^. In stony corals ^4,23–27^, a subset of these proteins comprises a conserved “biomineralization toolkit” whose evolution has been a recent focus of study ^4^, hinting at shared molecular strategies for calcification across diverse coral lineages. Similar to other biomineralizing invertebrates, the greatest emphasis rests on the highly acidic proteins - termed coral acid-rich proteins (CARPs) and skeletal/secreted aspartic acid-rich proteins (SAARPs) - and carbonic anhydrases ^28,29^. Highly acidic proteins are enriched in aspartic (Asp/D) and/or glutamic (Glu/E) acids (usually considered to be >35%), which in some cases form D/E stretches throughout the protein sequence, and are a feature seen to be shared by most – if not all – mineralizing phyla ^30,31^. The negatively charged sites interact strongly with cations, such as Ca^2+^, with ∼2-3 acidic residues coordinating each Ca^2+^ ion in the case of CARPs/SAARPs ^32^. Due to this, biomineralizing proteins frequently have the ability to coordinate Ca^2+^ reversibly, making acidic proteins central to classical hypotheses on carbonate biomineralization ^30,33^. In addition, stretches of negatively charged residues at biological pH prevent stable folding due to electrostatic repulsion between like-charged residues, which classifies them as intrinsically disordered proteins ^34^. Their intrinsic disorder at physiological pH further suggests dynamic roles in stabilizing mineral precursors and coordinating ions. Carbonic anhydrase complements these functions by catalyzing the hydration of CO_2_ to bicarbonate (HCO_3_^-^), which can be protein-coordinated ^32^ or, at elevated pH, converted to carbonate (CO_3_^2-^) ^35^ for CaCO_3_ precipitation, linking cellular metabolism to carbonate chemistry within calcifying compartments.

Beyond highly acidic proteins and carbonic anhydrases, homology of conserved sequences has enabled the identification of additional SOM proteins, including adhesion and structural proteins such as mucin and collagen, respectively ^24–27,36^. For other enigmatic proteins, functional information is hinted at from topological features such as the di-cysteine repeat in galaxin ^37^, although true function remains to be established. This gap has historically been filled using recombinant SOM proteins and crystal growth experiments ^32^ or NMR spectroscopy and X-ray crystallography for protein structure determination ^38^, which is expensive and time consuming. Alternatively, advances in AI-based structural predictions may allow examination of potential enzymatic active sites or the potential to bind or coordinate mineral ion precursors ^39^, that may direct future experiments ^40^.

A major unresolved question is how SOM proteins and mineral precursors are delivered from calicoblastic cells to the ECM in a controlled and coordinated manner. While classical secretion pathways account for a subset of skeletal proteins, increasing evidence points to vesicle-mediated transport ^17^, cytoskeletal trafficking ^5^, and dynamic recycling of calcifying compartments ^41^. How these pathways integrate to sustain continuous skeleton growth, particularly under extreme environmental conditions, remains poorly understood.

Photosymbiotic corals add further complexity as their endosymbionts (Symbiodiniaceae; ^42^) can enhance calcification by: (a) translocating photosynthate that supports ion transport and SOM synthesis ^43^, (b) removing host waste products (e.g., phosphate) that can inhibit carbonate nucleation ^44^, and (c) increasing CaCO_3_ saturation via CO_2_ uptake and an oxygen-rich microenvironment ^45^. Disentangling host-driven biomineralization mechanisms from symbiont-derived effects therefore requires model systems in which calcification occurs in the absence of photosymbiosis.

Here, we address these gaps by characterizing the SOM proteome of the asymbiotic, cold-water coral *Desmophyllum pertusum* (formerly *Lophelia pertusa*; Addamo et al., 2016), which maintains calcification under extremely unfavorable conditions of seawater pH down to ∼7.68 and aragonite saturation states of 1.31 ^46^. We then compared this proteome with published skeletal proteomes from the obligately photosymbiotic tropical coral *Stylophora pistillata* and the facultatively photosymbiotic temperate coral *Oculina patagonica* to evaluate how photosymbiosis may affect biomineralization ^27,47^. These three taxa are all colonial scleractinians (Figure 1), enabling functional comparisons across shared skeletal growth modes while spanning distinct trophic states and habitats. Boron isotope (δ^11^B) measurements of coral skeletons indicate that asymbiotic cold-water corals strongly up-regulate the pH of their ECM, elevating it by approximately 0.8-1.0 units (ΔpH) above ambient seawater, substantially more than the 0.4-0.5 unit elevation typically observed in shallow-water species ^48^. As such, it has been hypothesized that the composition and properties of the SOM may contribute to the variability in ΔpH observed among scleractinian corals ^49^. By leveraging this symbiont-free model and integrating comparative proteomics, genomics, and AI-based structural prediction, we assess whether coral calcification is underpinned by a conserved, dynamic biomineralization toolkit that operates across contrasting trophic modes and environmental regimes.

**Figure 1.**
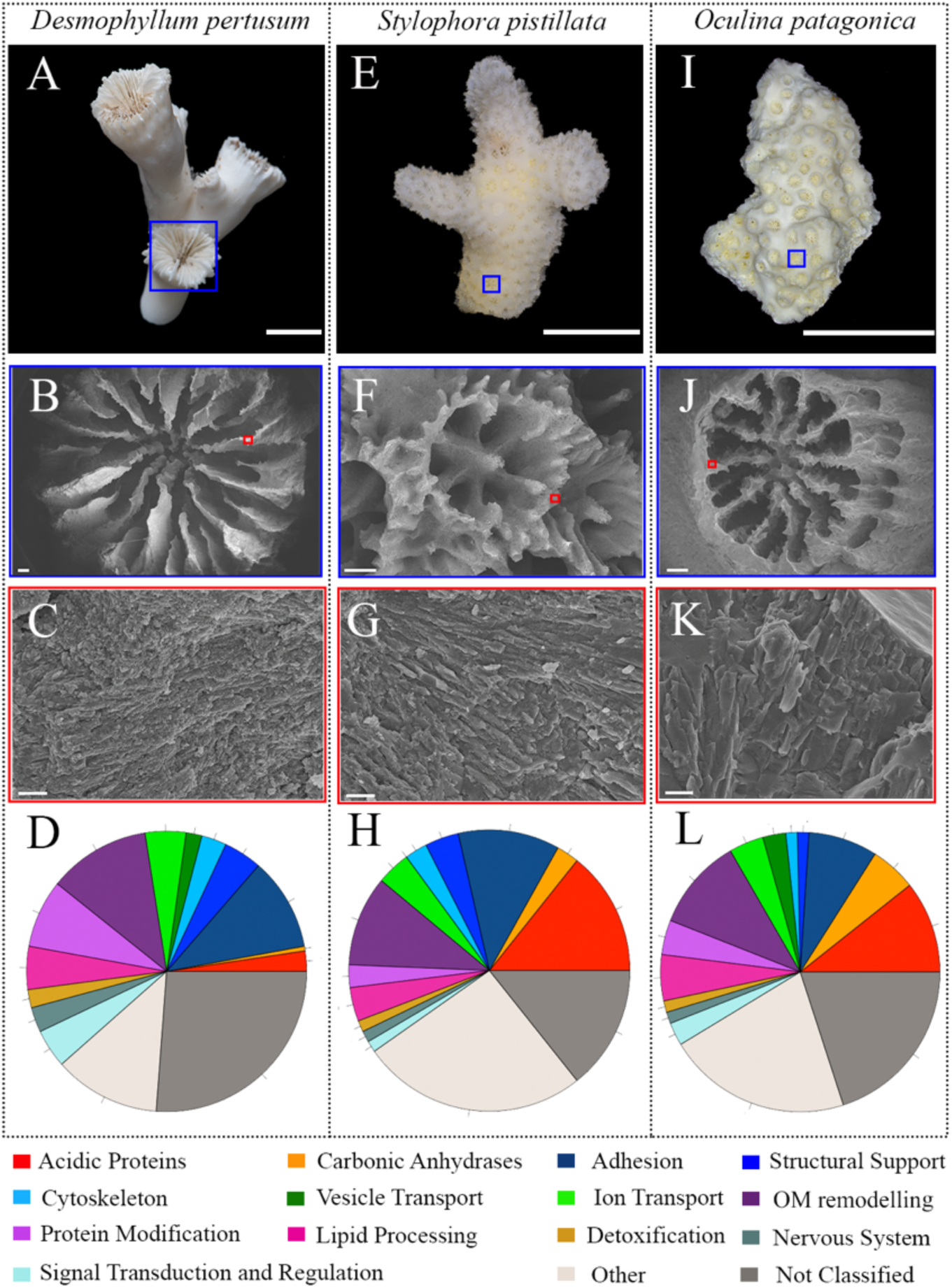
Macrographs of skeletal fragments, SEM images, and SOM protein function classifications. in: (A-D) *D. pertusum* (n = 382), (E-H) *S. pistillata* (n = 77), and (I-L) *O. patagonica* (n = 71). B, F, and J are SEM images of the regions indicated by blue boxes in A, E, and I, respectively. C, G, and K are higher-magnification views of the regions indicated by red boxes in B, F, and J, respectively. Protein functions are based on gene ontology and annotation. Similar functions are colored similarly (i.e., physical framing of the calcifying space in shades of blue for Adhesion, Structural Support, and Cytoskeleton; transport pathways in shades of green for Vesicle Mediated Transport and Ion Transport; adjusting the organic composition of the calcifying space in shades of purple for OM Remodeling, Protein Modification, and Lipid Processing; and signaling in shades of cyan for Nervous System and Signal Transduction and Regulation). Scale bars on first, second, and third rows are 1 cm, 200 µm, and 1 µm, respectively.

## Results and Discussion

### Shared biomineralization toolkit across coral taxa from shallow to deep-sea environments

Three hundred and eighty-two SOM proteins predicted from the *D. pertusum* genome assembly were detected by LC-MS/MS analysis (**Table 1**; Data S1). These proteins were then grouped by their proposed function based on amino acid composition, gene ontology analysis, and annotation and compared with previously-published skeletal proteomes from *S. pistillata* and *O. patagonica* ^24,47^. Despite significant differences in skeletal micro-density between *S. pistillata* (2.2 ± 0.07 mg/mm^3^), *O. patagonica* (2.5 ± 0.09 mg/mm^3^), and *D. pertusum* (2.6 ± 0.09 mg/mm^3^) (One-way ANOVA, *F_2,9_* = 12.045, *p* = 0.005), we observed similar: 1) skeletal structural patterns (Figure 1B,E,H), 2) distributions of protein functions (Figure 1C,F,I), 3) intraskeletal water and OM content of 2.9-3.1% by weight (Figure S1), as found in other shallow and cold-water corals ^50,51^, and 4) amino acid composition biased toward Asx and Glx (Table S1), in line with previous studies ^52,53^. Our OrthoFinder analysis revealed (Figure 2): 1) a core set of orthogroups – clusters of genes derived from a common ancestor – shared by all three species within the same functional groups; 2) at least one orthogroup per functional group (except for the ‘Nervous System’ group) shared between the cold-water coral and one of the two shallow-water corals, suggesting a conserved toolkit; and 3) orthogroups unique to *D. pertusum*. Overall, ∼16% of SOM protein isoforms in the cold-water coral fall within an orthogroup shared with at least one of the two shallow-water coral species.

**Figure 2.**
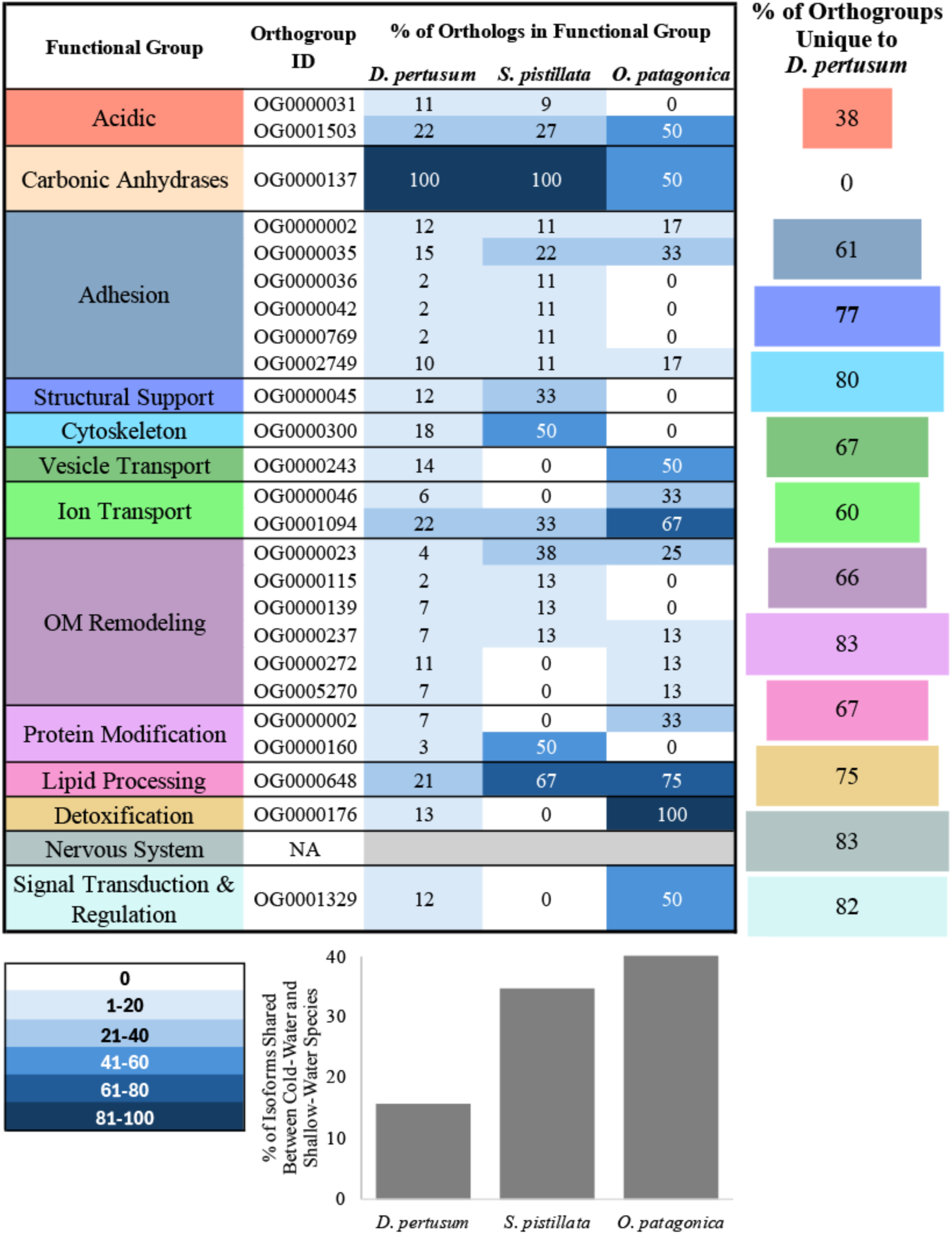
OrthoFinder analysis of SOM protein isoforms across the cold-water coral *D. pertusum* and the shallow-water corals *S. pistillata* and *O. patagonica*. In the heat-map (top), for each functional group, values indicate the percentage of orthologs assigned to each orthogroup per species, with columns labelled by species (*D. pertusum*, *S. pistillata*, *O. patagonica*). Cell color reflects that percentage according to the heat-map scale (bottom left): white = 0, with increasingly dark blue shading in bins of 1–20, 21–40, 41–60, 61–80, and 81–100. ’NA’ (grey shading) denotes functional groups with no assigned orthogroup shared between taxa. The horizontal bar chart reported on the right shows the percentage of orthogroups within each functional group unique to *D. pertusum*. The vertical bar chart (bottom right) reports, per species, the percentage of SOM protein isoforms that fall within an orthogroup shared between the cold-water and shallow-water corals.

**Table 1.**
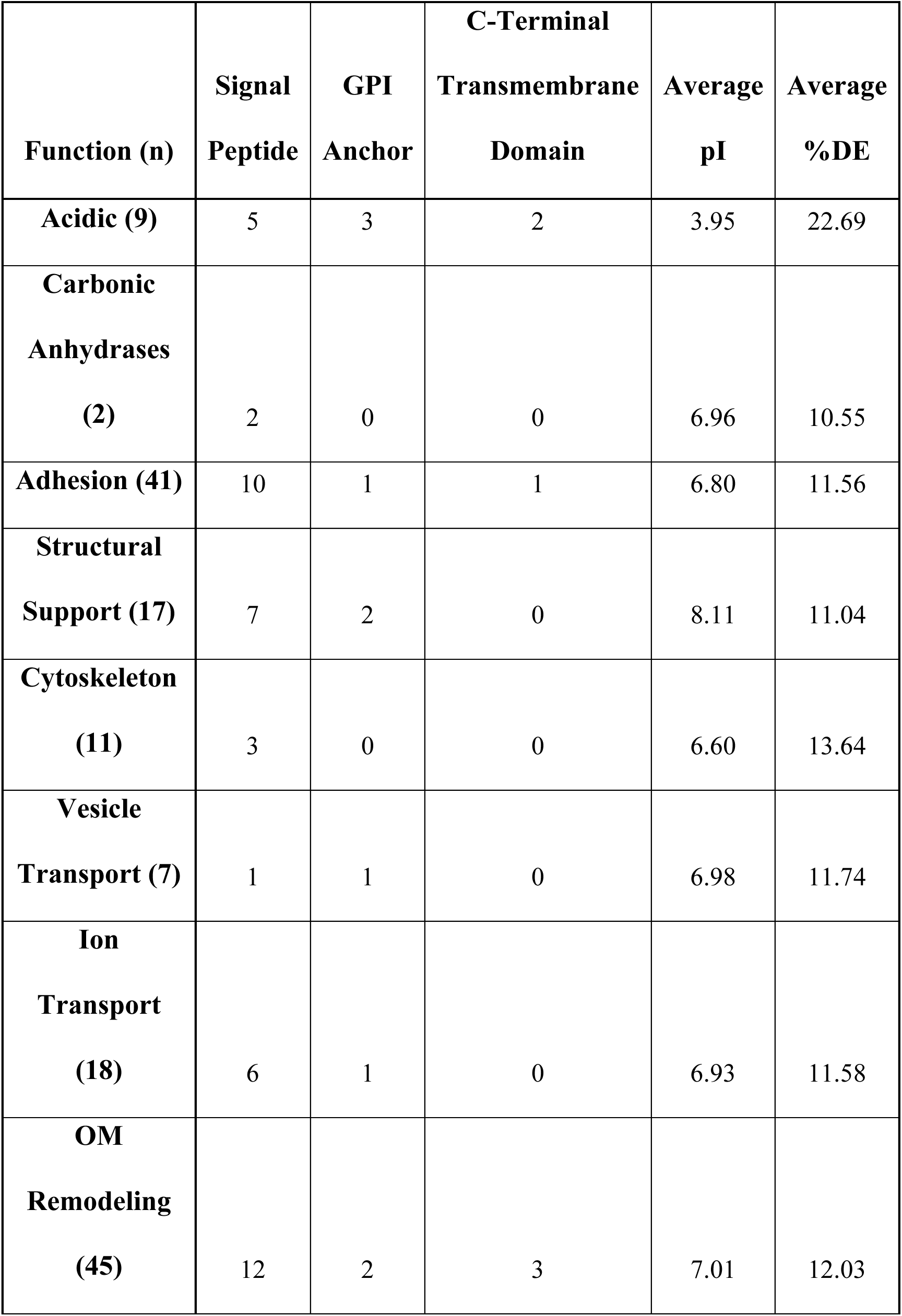
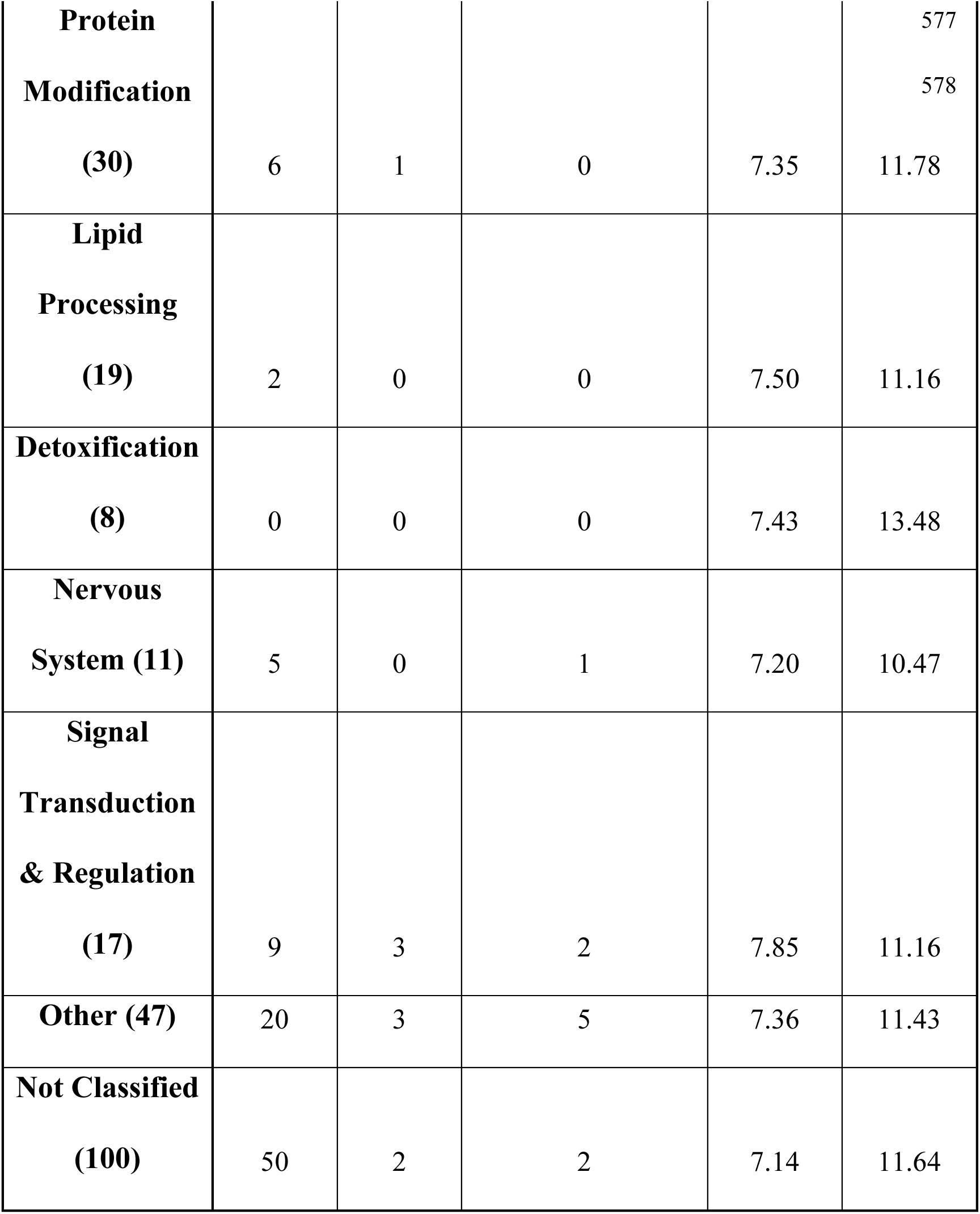
*D. pertusum* skeletal proteins. Summary of 382 identified proteins in *D. pertusum* skeleton grouped by proposed function.

This preliminary comparison of three species reveals significant overlaps in biomineralization machinery across taxa. We therefore feel confident exploring the SOM proteins of *D. pertusum*, the builder of extensive deep-sea reefs without the aid (and complications) of photosymbionts, as a novel model system to expand the working biomineralization model originally built exclusively from shallow-water coral species.

### Calicoblastic cell layer structure and signaling

Like other studies ^24–27,32,36,47,52^, we observed in *D. pertusum* SOM a variety of proteins that likely provide structure to the ECM. We detected 42 calcium-dependent adhesion cadherins, integrins, contactin-associated proteins, mucins, hemicentins, coadhesins, and similar adhesion proteins. These have been suggested to (i) constitute an extracellular matrix that adheres to the newly formed skeleton and (ii) attach calicoblastic cells to this skeleton-blanketing matrix in corals ^24,25,27^ and molluscs ^54–56^. Many of these adhesion proteins contain thrombospondin type 1 repeats, von Willebrand factor type A and epidermal growth factor-like domains, which are typical of ECM proteins involved in cell–cell and cell–substrate adhesion and in the binding of other macromolecules ^57^. We also detected several fibrillar-type proteins, including nine annotated as collagens, which are common in the extracellular matrix of vertebrate bone and in mineralized structures of other non-scleractinian cnidarians ^58^ and scleractinian coral skeletons ^24,25,27,36^. In addition, we identified proteins involved in collagen fibril organization (hemagglutinin/amebocyte aggregation factor-like, Deleted in malignant brain tumors 1 protein) and collagen trimmers (tolloid-like protein 1 isoform X2, Fibrinogen beta chain). In particular, the high redundancy in skeletal collagens supports previous gene expression-based evidence for multiple types of calicoblastic cells ^59^, consistent with earlier geochemical work suggesting that distinct calicoblastic cell types may drive mineral growth in different skeletal regions ^7^.

*D. pertusum* also contains two galaxin-like proteins, which were first identified in the exoskeleton of the scleractinian coral *Galaxea fascicularis* ^37^ and observed in the exoskeleton of other stony corals ^26^. Although its function remains elusive, galaxin is associated with post-larval onset of coral calcification ^60^ and has been proposed as a potential collagen binding protein due to shared di-cysteine similarity with usherin ^61^. All three stony coral taxa considered in the current study retain in their skeleton roughly the same proportion of SOM proteins assigned to nervous system-like functions. Of 11 *D. pertusum* SOM proteins with GO terms or annotation related to the nervous system, six are related to adhesion (i.e., neural cell adhesion molecules and neuroligins). Their function in biomineralization remains unclear.

Signaling between the calicoblastic cell membrane and the ECM is crucial for controlling mineralization. We found 17 *D. pertusum* proteins associated with signal transduction. These include a bone morphogenetic protein (BMP) and multiple proteins associated with the Wnt signaling pathway, among others. BMPs and core Wnt-pathway genes have been previously linked to biomineralization ^62^. BMPs are expressed in the calcifying epithelium of multiple coral species ^63^ and may respond to changes in Ca^2+^ within the ECM to promote differentiation of distinct calicoblastic cell types ^59^.

### Multiple mineral and biomolecule export pathways from the cell

Of the 382 proteins sequenced from *D. pertusum* skeleton, 189 proteins (ca. 50%) exhibit documented ‘classical’ features designating them for export (Pathway 1 in **Figure 3**), which is similar to the percentage found by ^27^. Specifically, 138 (36%) are predicted to contain a signal peptide (Table 1; Data S1). Of these, 10 are also predicted to possess a C-terminal GPI anchor for deposition at the plasma membrane, which requires the presence of an N-terminal signal peptide ^64^, 16 contain a transmembrane span within 30 amino acids of the C-terminus - a tail anchor - suggesting that they are embedded within the cell membrane with the N-terminal end oriented toward the ECM ^65^, and 59 contain non-signal/GPI/tail-anchor transmembrane spans. Post-translational modifications may also aid in protein export; we found acyl-protein thioesterase 1-like and palmitoyl-protein thioesterase 1-like which modify proteins that lack a signal peptide for transport to the plasma membrane ^66^, as well as palmitoyl hydrolases which remove the PTM extracellularly.

**Figure 3.**
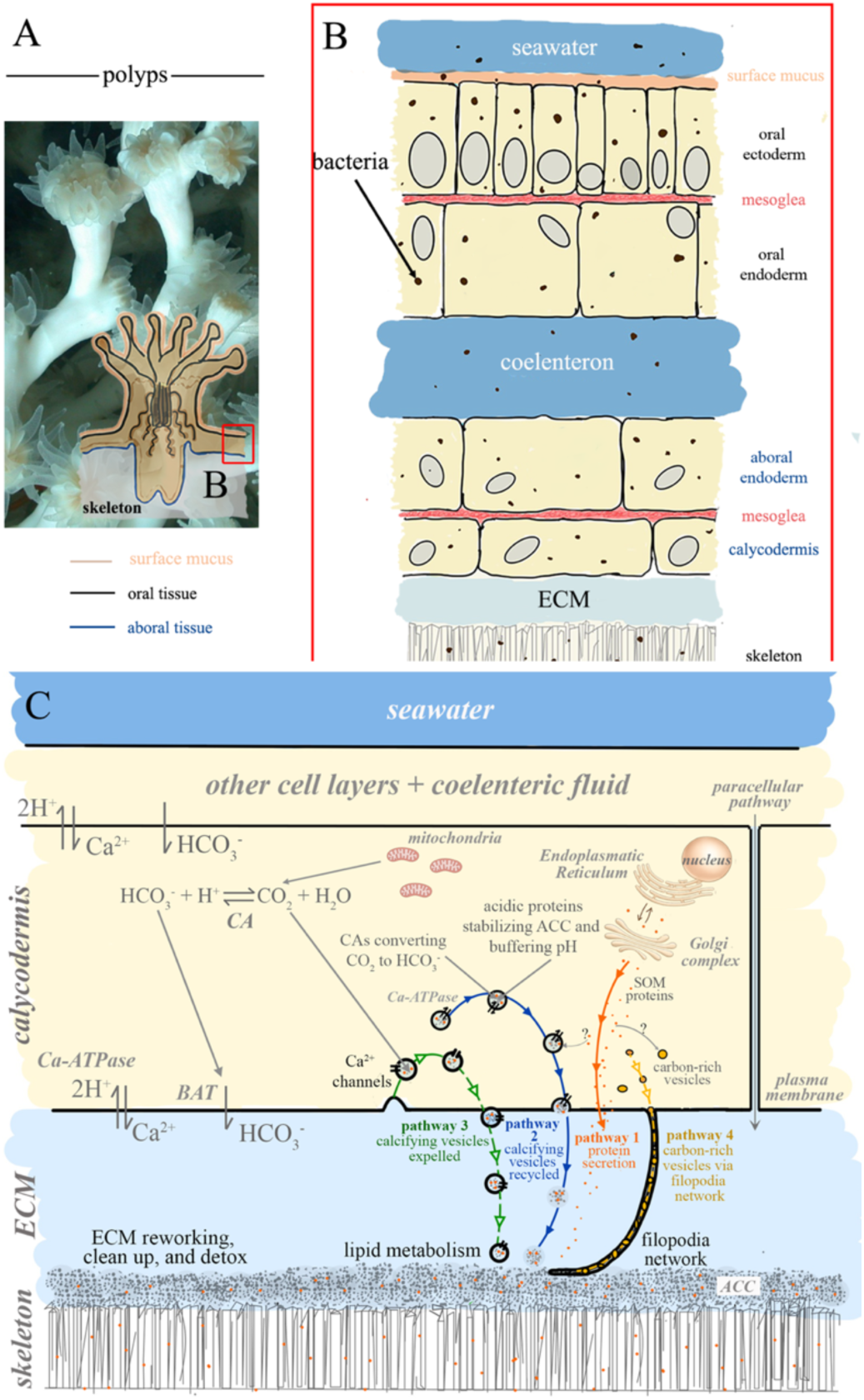
Structure of a coral polyp and processes involved in biomineralization. (**A**) Close up of *Desmophyllum pertusum* polyps with overlaid drawing of a coral polyp highlighting surface mucus, the oral tissue, and the aboral tissue. (**B**) Schematic representation of a polyp’s cell layers (cross-section) (59) and location of bacteria within a polyp (112). (**C**) Four pathways deliver material from the calicoblastic cell to the growing skeleton. Pathway 1 (protein secretion) and Pathway 2 (macropinocytosis-derived calcifying vesicles recycling within the plasma membrane) are established pathways (solid lines). Pathway 3 and Pathway 4 are the two pathways proposed in this study (dashed lines). In Pathway 3, calcifying vesicles are fully exocytosed into the ECM. In Pathway 4, carbon-rich vesicles are transported through the filopodia network directly into the growing skeleton. Within the calcifying vesicles, carbonic anhydrases catalyze the conversion of CO_2_ to HCO_3_^-^, while acidic proteins stabilize amorphous calcium carbonate (ACC) and sequester protons, helping buffer pH. Text in black refers to functional categories of proteins that were identified in the current study and are included for the first time in a coral biomineralization model (e.g., ECM reworking, clean up, and detox, lipid metabolism). Cellular compartments (not represented according to their relative sizes) and proteins that were not specifically identified in this study are shown in gray font.

For the remaining half of *D. pertusum* SOM proteins, assigned functions strongly support the role of vesicular and cytoskeletal transport mechanisms. In calicoblastic cells, mitochondria are abundant ^15^ and supply much of the CO_2_ used for calcification ^67^. Respired CO_2_ is converted to HCO_3_^-^ by a cytosolic pool of carbonic anhydrases (CAs) ^68^ and delivered to the ECM via plasma membrane bicarbonate transporters (BATs) ^16,69^. Within the ECM, elevated pH, maintained by Ca-ATPases that both supply Ca^2+^ and extrude H^+^, shifts carbonate chemistry toward CO_3_^2-^ and promotes CaCO_3_ precipitation ^70^. Studies have shown that the ECM can be endocytosed by macropinocytosis, a non-clathrin-dependent, actin-driven process, into calicoblastic cell vesicles 100s nm in size ^13^, consistent with similarly sized vesicles observed in earlier microscopy work ^71^. Abundant vesicular transport across the calicoblastic cell layer, including vesicle fusion with the plasma membrane, has been documented in *Acropora yongei* ^15^, *A. hebes* ^72^, and *Pocillopora damicornis* ^73^. Notably, we identified a Rab11 homolog (ras-related protein Rab-11A), a key regulator of slow endosomal recycling ^74^. In *S. pistillata*, Rab11 is expressed in the epidermis ^59^ and was recently shown to associate with macropinosomes formed by ECM endocytosis at the calicoblastic apical membrane, promoting recycling back to the plasma membrane ^17^. Such vesicles, hereafter termed “calcifying vesicles”, may either take up SOM proteins present in the ECM or incorporate them within calcifying cells via an as-yet-unknown mechanism. These vesicles may also promote the formation of amorphous calcium carbonate (ACC) particles, hundreds of nanometers in diameter, which have indeed been observed in calicoblasts. ^5,14,17^. In our dataset, we also detected multiple proteins linked to voltage-gated Ca^2+^ channel activity (five VWFA and cache domain-containing protein 1-like, polycystic kidney disease 1-related protein, polycystic kidney disease protein 1-like 2, ZP domain-containing protein-like, Sodium/potassium/calcium exchanger 5) together with polycystins known to modulate Ca^2+^ channels ^75^. This aligns with prior work supporting active transcellular Ca^2+^ transport in coral calcification ^47,76^. These proteins may support Ca^2+^ loading from the cytosol or mitochondria into the calcifying vesicles ^77^, which are then exocytosed into the ECM, thus explaining their presence in the skeleton (Figure 3).

Building on this, a subset of calcifying vesicles reaches the plasma membrane, expels its contents (e.g., CAs, acidic proteins, ACC) into the ECM, and fuses with the plasma membrane, thereby recycling vesicle membrane and associated transport machinery, including BATs and Ca-ATPases (Pathway 2 in Figure 3). Such recycling would help explain why certain membrane components (transporters and Ca-ATPases) are absent from skeletal proteomes. In parallel, the finding of SOM vesicles that lack Rab11 ^17^ supports our hypothesis that an additional vesicle population exists - potentially enriched in Ca^2+^ channels identified in the current study - that is not recycled but instead are fully exocytosed into the ECM (Pathway 3 in Figure 3).

Actin and myosin are important for the cytoskeletal train of export, often found in cellular elements such as filopodia, which have been observed extensively in calicoblastic cells ^5,78,79^. Shapes of calicoblasts, particularly desmocytes, are known to conform to those of the underlying skeleton ^6^, enabling cellular projections into the skeleton/calcifying space ^80–82^ and allowing expansion and contraction of the calcifying space on timescales of minutes ^5^. In *D. pertusum* SOM, we identified two actins (actin cytoplasmic-like and actin cytoplasmic) and several proteins involved in actin binding (radixin-like, plastin-3 like isoform X1, myosin light chain kinase, smooth muscle-like, gelsolin-like protein 2 and 1, coronin-6 like, and coactosin-like protein), which along with the myosins and gelsolins, could support size and shape flexibility within the calicoblastic space during aragonite crystal growth ^24^. The proteins in this cytoskeletal network, including calsyntenins ^83^, may also allow the transport of vesicles containing proteins that lack traditional signaling components ^84,85^ along microtubules ^86^ for eventual exosomal release ^87^ and budding ^88^. These proteins may therefore provide the first biochemical evidence of the carbon-rich vesicles observed in calicoblastic filopodia by ^5^. Accordingly, we propose that proteins are packaged into carbon-rich vesicles and transported along the cytoskeletal network via filopodia that extend into the ECM and the growing skeleton, where they are subsequently released through exosomal secretion into the calcifying space (Pathway 4 in Figure 3). Moreover, the fact that we find cytoskeletal proteins in *D. pertusum* suggests that this species also possesses filopodia projections, which have not yet been documented histologically in cold-water corals.

The possible role of vesicular pathways in coral biomineralization is also supported by the presence of SOM proteins involved in lipid metabolism found in all three taxa considered in the current study. Lipid binding and processing is crucial for recycling exocytosed vesicles containing SOM biomolecules and mineral precursors. Proteins containing low-density lipoprotein receptors (LDLR) play a crucial role in regulating lipid metabolism ^89^. LDLR have been observed in invertebrate minerals from corals ^26^ to sea urchins ^90–92^ and their deficiency results in reduced bone mineral density ^93^. Additionally, the *D. pertusum* SOM contains eight lipoxygenases which oxidize polyunsaturated fatty acids, and SOMs from all comparison taxa present vitellogenin, a lipid carrying protein ^94^.

### A Dynamic Extracellular Matrix

Stony coral SOM proteins contain several potential sites of post translational modifications (PTMs), mainly glycosylation (i.e., the covalent attachment of single sugars or glycans to select residues of target proteins) and phosphorylation (i.e., the reversible addition of a phosphate group to specific amino acids, typically serine, threonine, or tyrosine). Glycosylation is a major form of PTM which expands the functional diversity and structural heterogeneity of proteins, characterizing 50-70% of human proteins ^95,96^. Glycosylation has a significant impact on protein-mediated biomineralization in general (e.g., nucleation, crystal growth, and matrix assembly) ^97^ and on protein function, in particular, being directly related to protein solubility, folding, stability, activity protection from proteases, and subcellular targeting. Proper protein glycosylation is also important for the formation of protein complexes and for modulating protein–protein interactions ^98^. Glycosylation of coral SOM proteins has been documented in several scleractinian species and is thought to contribute to skeletal formation ^29,99,100^. It is therefore not surprising that we found polypeptide N-α-acetylgalactosaminyltransferases (ppGalNAcTs) (polypeptide N-acetylgalactosaminyltransferase 1-like, polypeptide N-acetylgalactosaminyltransferase 2-like) which are able to produce densely glycosylated mucin glycoproteins ^101^.

We also detected several proteins involved in the synthesis of polysaccharides or glycoproteins, specifically two glycosyltransferases (beta-1,4-N-acetylgalactosaminyltransferase 3-like) ^102,103^, and in glycan degradation (Tissue alpha-L-fucosidase [*Desmophyllum pertusum*] ^104^. Extracellular glycotransferases may be limited by access to nucleoside sugars ^105^, although sugar donor release by platelets has been observed ^106^. Interestingly, *D. pertusum* SOM contains nucleoside phosphatases and diphosphatase that may alleviate this limitation. We also found three β-galactosidases (two beta-galactosidase-1-like protein 2 and one beta-galactosidase-like), which are enzymes that remove β-linked terminal galactosyl residues from substrates such as glycoproteins and glycosaminoglycans ^107^.

Protein phosphorylation provides a mechanism for the rapid, reversible control of protein function. Target proteins are phosphorylated at specific sites by one or more protein kinases (PKs) ^108^, and these phosphates are removed by specific protein phosphatases (PPs) ^109^. We detect several PKs (protein kinase C-binding protein NELL2-like, protein O-mannose kinase-like, protein amalgam-like, inactive tyrosine-protein kinase transmembrane receptor ROR1-like, neural cell adhesion molecule 1-B-like, titin-like isoform X1 and X6) and PPs (inositol monophosphatase 3-like, two receptor-type tyrosine-protein phosphatases, two acid phosphatase type 7-like). Phosphorylation adds negative charge to amino acid side chains, and negatively charged amino acids (Asp/Glu) can sometimes mimic the phosphorylated state of a protein ^110^, temporarily stabilize ACC or nucleate the mineral under appropriate conditions ^30^. Stabilization of ACC may occur via phosphate rich organic matrix proteins and by single phosphoamino acids ^111^.

Once enzymes have completed their role in mineral formation from the ECM, they may become a liability upon degradation or due to malformation. The endosomal/lysosomal pathway is viewed as a means of containing, transporting, and degrading unwanted materials due to toxicity or damage ^112,113^. We found several *D. pertusum* SOM proteins associated with endosomes and lysosomes – such as ras-related proteins, Rab-GDP, cubilin-like proteins - supporting this pathway. Furthermore, peptidases observed in coral SOM may also play a role in removing proteins that are no longer needed or potentially harmful (e.g., any malformed proteins) ^114^, while the observed peptidase inhibitors could in turn help maintain a balance between protein degradation and maintenance. *D. pertusum* SOM contains >10 peptidases. We identified two metalloproteinases (zinc metalloproteinase nas-15-like isoform X2, zinc metalloproteinase nas-39-like), enzymes capable of cleaving extracellular matrix proteins and have been suggested to play a key role in ECM remodeling ^41^.

The byproducts of organic matrix modification and degradation include oxidative stress molecules such as hydroperoxide and peroxinitrite substrates ^115^. We detected proteins involved in detoxification by neutralizing these compounds in *D. pertusum* SOM as well as in the comparison taxa. These include peroxidasin, thioredoxin, glutathione transferases, and peroxiredoxins. Cnidarians contain representatives from three subfamilies of peroxiredoxins which bind to and neutralize peroxides ^116^, and in mammals are involved in membrane repair following oxidation ^117^. Hence, it appears that stony coral SOM contains a repertoire of proteins not just for extracellular matrix reworking, but also for its later clean up.

### Acidic Proteins and Carbonic Anhydrases

‘Highly acidic’ proteins in coral skeletal research have been defined using 35% D+E as a lower bound (e.g., in the first coral genome by ^118^). Although the abundance of aspartic and glutamic amino acids across ‘standard proteins’ has been noted as 5.3 and 6.2%, respectively ^30^, no lower bound for generally ‘acidic’ coral skeletal proteins has yet been proposed. To query the acidic SOM proteins of *D. pertusum*, we therefore searched for proteins that display >14.5% D+E and a pI <4.5 ^30^, analyzed using Multiple Protein Profiler 1.0 ^119^. Seven CARPs/SAARPs have been characterized to date: CARP1, CARP2, CARP3, CARP4/SAARP1, CARP5/SAARP2, CARP6, acidic SOMP/SAARP3. In *D. pertusum* we detected CARP4/SAARP1 and acidic SOMP/SAARP3 and seven novel SAARPs that we named SAARP7-13 (Data S1).

Negatively charged Asp/Glu (D/E)-rich regions in acidic proteins ^47,120^ have been proposed to bind free Ca^2+^, promoting a Lewis acid–base reaction that displaces a proton from HCO_3_^-^, facilitates its conversion to CO ^2-^ and thereby favors CaCO precipitation ^32^, likely as ACC within calcifying vesicles ^5,14,121,122^. However, specific Ca^2+^ binding or coordination functions have not been assigned to individual acidic proteins beyond CARP3 and CARP4/SAARP1 in *S. pistillata*. Given the diversity of sequence motifs among proteins classified as “acidic” across taxa, this group likely encompasses a range of still-undefined roles in mineral formation. We propose that one such role is transient pH buffering within calcifying vesicles during proton sequestration following HCO_3_^-^ deprotonation, helping sustain ACC formation. This mechanism could help explain the recurrent detection of acidic proteins and absence of proton pumps and channels across all coral skeletal proteomes published to date. Bicarbonate may already be present in calcifying vesicles through endocytosis of ECM (as described above), or it may form within the vesicles when diffusing CO_2_ is hydrated to HCO_3_^-^ by CAs ^123^, which we detected in *D. pertusum* in the present study (active α carbonic anhydrase type 2-like and an inactive α carbonic anhydrase type 2-like) and which have been found in multiple other stony coral SOMs ^24,25,27,36,47^.

### Describing protein functions through structure prediction and analysis

Inferring protein functions from primary sequence alone can be challenging, as several SOM proteins show little sequence homology across taxa outside Scleractinia ^4^. Here we integrated sequence homology with AI-based structural predictions on CARPs/SAARPs and carbonic anhydrases using AlphaFold 3 (AF3) ^124^ to predict the three-dimensional models of the sequences and consequently analyze their properties. Additional structural predictions were performed on a subset of species-specific proteins described in detail in the Supplemental section (Figures S5-S8).

The CARP4/SAARP1 family is one of the most highly detected proteins in scleractinian corals ^24,27,36,47^. Our structural analysis revealed that both CARP4/SAARP1 and acidic SOMP/SAARP3 in *D. pertusum* (FUN_020759-T1, FUN_016248-T1) are predicted to fold into a β-sandwich topology of 5- and 7-stranded antiparallel sheets with high resemblance to Ig or fibronectin type III folds (Figure 4A, Figure S2A) ^125^.

**Figure 4.**
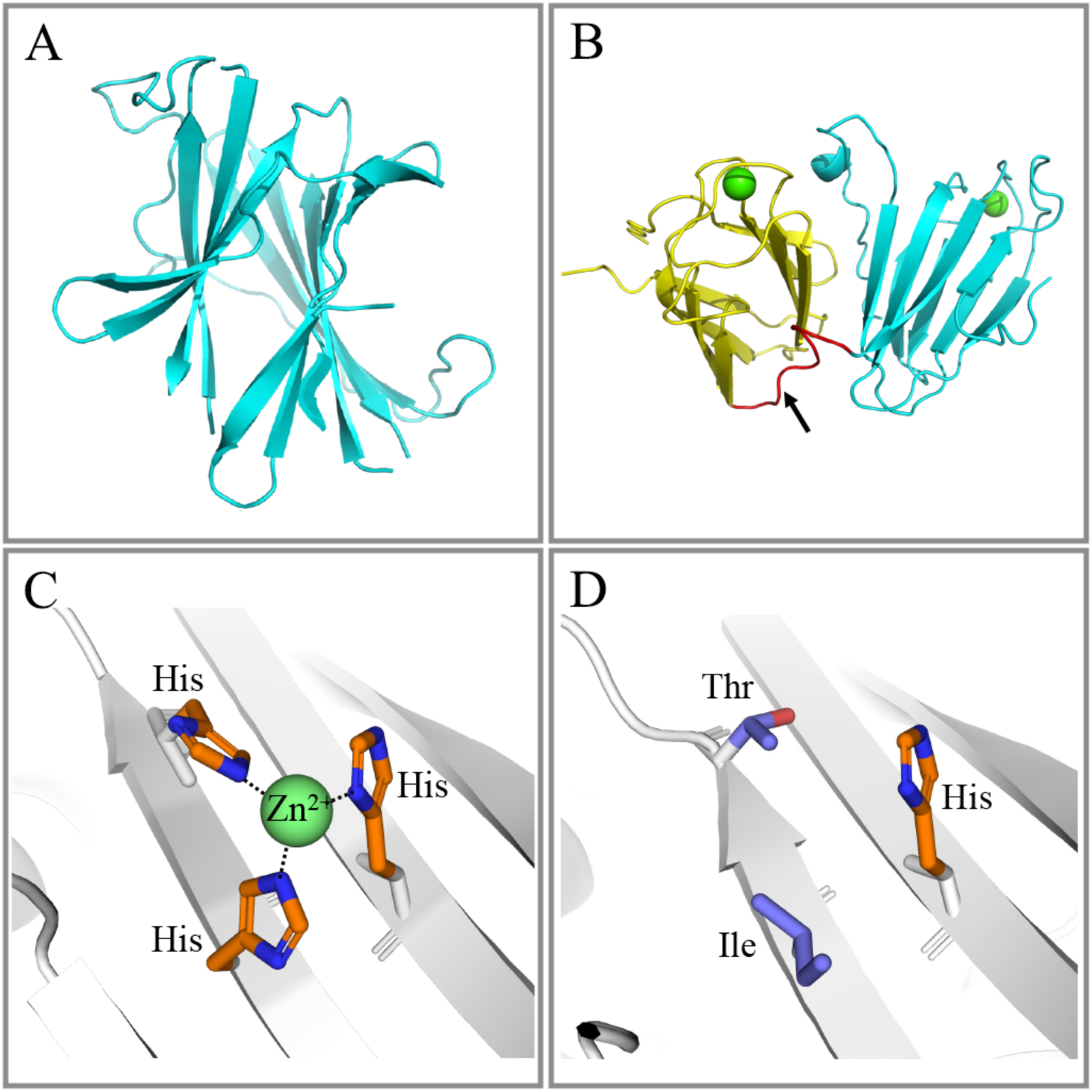
Structural predictions of representative acidic proteins and carbonic anhydrases from *D. pertusum* skeleton. (**A**) CARP4/SAARP1 model generated by AF3 shows the core of the prediction (the undefined D/E rich segment was removed for clarity). The model exhibits the topology of a β-sandwich consisting of 5 and 7 strands resembling an Ig fold, in which flat arrows represent β-sheets. (**B**) The core of the two–β-topology (shown in cyan and yellow) connected by a short linker (red) indicated by a black arrow, found in SAARP12 from *D. pertusum*. At the edge of each domain there is a Ca^2+^ (green spheres) binding site. (**C**) Active α carbonic anhydrase type 2-like protein from *D. pertusum* clearly showing Zn^2+^ is coordinated by 3 His residues rendering the enzyme active. (**D**) Inactive α carbonic anhydrase type 2-like protein from *D. pertusum* clearly showing the lack of two essential His residues that are replaced by Ile and Thr, now incapable of coordinating Zn^2+^, resulting in an inactive biomolecule.

SAARP12 (FUN_024181-T1) is a newly identified acidic protein in *D. pertusum.* This protein exhibits a topology made of two β-sandwiches connected by a short linker (12 residues) (Figure 4B, Figure S2B). The same topology was found in a candidate acidic protein that had been overlooked by sequence homology-based annotation in *S. pistillata*: tolloid-like protein 2 (g5735.t1, NCBI accession XP_022799127.1; ^36^). Each of the domains contain a putative Ca^2+^ binding region assessed by DALI search for similar topologies available in the PDB ^126^. In this context, there are similar folds in the PDB only for each of the β-sandwiches, but no topology for two of them in tandem. For each of them, the similarities include: neuropilin-2 (PDB entry: 2qqo-A) and tumor necrosis factor-inducible gene 6 protein (PDB entry: 2wno-A), mannan-binding lectin serine protease 1 (PDB entry: 5ckn-D) all containing a Ca^2+^ ion at each end of the β structure with similar coordinating residues. In all AF3-predicted models, some segments lack a defined topology and are predicted to be disordered or to adopt multiple conformations. Given the known limitations of AF in these cases, we infer that when these proteins (including CARPs, SAARPs, and others) encounter Ca^2+^ ions in seawater, their intrinsically disordered D/E-rich regions may undergo conformational compaction stabilized by specific, chelation-like ionic interactions with acidic side chains. This has been demonstrated experimentally for the coral acid-rich protein AGARP from *Acropora millepora*, which exhibits a valency-sensitive electrostatic collapse driven by Ca^2+^ and Mg^2+^ binding to its Asp/Glu carboxylate groups ^127^. In line with this, and as previously suggested ^32^, we further propose that these D/E-rich regions contribute Ca^2+^ ions to the calcification process.

We detected an active α carbonic anhydrase type 2-like (FUN_012693-T1) in *D. pertusum*, displaying a canonical CA fold with the three-histidine coordination of Zn^2+^ in the active site (Figure 4C). AF3 also revealed a catalytically inactive (‘silent’) CA-related protein bearing an acidic segment (FUN_020142-T1), despite this protein carrying the same α carbonic anhydrase type 2-like annotation on the basis of sequence homology. This protein displays the structural characteristics typical of CAs but lacks Zn^2+^ binding capability, as two of the histidine residues are substituted by Thr and Ile and are not compensated for upon folding, rendering the molecule enzymatically inactive (Figure 4D). It seems that these inactive (“silent”) forms are not catalytically dead ends - they are repurposed signaling and scaffolding proteins that maintain the CA fold for protein–protein interactions, Ca^2+^ signaling control, and cellular localization of active CAs ^128,129^. In this context, they have evolved to perform important structural, regulatory, and signaling roles. However, the role of the disabled CA in *D. pertusum* remains enigmatic.

### An integrated model of coral biomineralization

Our comparative analysis of the *D. pertusum* skeletal proteome against those of *S. pistillata* and *O. patagonica* shows that, despite stark differences in trophic mode and habitat, stony corals share a broadly conserved “biomineralization toolkit” that supports a dynamic, highly regulated calcification machinery. Our findings open a window into the ‘black box’ of stony coral biomineralization by addressing how proteins synthesized by calicoblastic cells are incorporated into the skeleton in a regulated fashion. Four non-mutually exclusive pathways likely contribute to the delivery of material from calicoblastic cells to the growing skeleton (Figure 3). Approximately half of coral SOM proteins possess canonical secretory signals - such as signal peptides, C-terminal anchors, and GPI anchors - enabling their export from calicoblasts via traditional secretory pathways (Pathway 1) ^130^. The remaining proteins may enter macropinocytosis-derived calcifying vesicles ^13^, within which ACC forms. Upon reaching the plasma membrane, these vesicles may either discharge their contents (e.g., CAs, acidic proteins, ACC) into the ECM and fuse with the plasma membrane ^15,17^, thereby potentially recycling both membrane and associated Ca-ATPases and BATs (Pathway 2), or be fully exocytosed into the ECM and incorporated into the growing skeleton (Pathway 3, proposed in this study). Alternatively, proteins may be packaged into carbon-rich vesicles ^5^ and transported along the cytoskeleton via filopodia that extend into the ECM and the growing skeleton, where they are subsequently released through exosomal secretion (Pathway 4, proposed in this study).

Pathways 2 and 3 may help resolve a long-standing conundrum regarding the origin and composition of ACC-bearing vesicles observed in coral tissues ^14,121^. ACC formation requires concurrent delivery of inorganic carbon (HCO_3_^-^ and/or CO_3_^2-^), Ca^2+^, and effective proton removal. Bicarbonate may enter calcifying vesicles through endocytosis of ECM, via diffusion of mitochondrial-derived CO_2_ into the vesicles followed by CA-catalyzed hydration to HCO_3_^-^, and/or via BATs embedded in the vesicle membranes. Ca^2+^, in turn, may be loaded via Ca^2+^ channels in vesicles destined for full exocytosis, or via Ca-ATPases retained in recycling vesicles, which could also support intra-vesicular pH regulation by exporting H^+^ out of the vesicle lumen. Coral primary cell cultures, which secrete ECM and precipitate aragonite into the overlying medium ^52,120,131^, offer a tractable system in which exported vesicles could be isolated directly from the calcifying medium and characterized, analogous to ultracentrifugation- and proteomics-based workflows already established for vertebrate matrix vesicles ^132,133^. Furthermore, we hypothesize that acidic proteins may facilitate ACC precipitation within the calcifying vesicles by chelating Ca^2+^ and transiently buffering protons released during mineral formation. Both functions are testable using recombinant proteins in *in vitro* precipitation assays, as previously demonstrated for *S. pistillata* ^32^. Within this framework, acidic proteins do not contribute to skeleton formation as a single functional class but as a diverse suite of molecules with distinct architectures and putative roles.

As the ECM is continuously supplied with mineral precursors and newly secreted proteins, it must be dynamically remodeled to sustain optimal calcification conditions. The functional analysis of SOMs from *D. pertusum* supports a model of extensive ECM reworking, in which proteins are altered by the addition and removal of extracellular post-translational modifications. We also identified different ways by which the large pools of proteins and lipids ^53^, likely delivered via vesicle export, can be degraded once they become obsolete or damaged. Finally, multiple pathways appear to contribute to ECM detoxification as these biomolecules are degraded.

By integrating proteomics, comparative genomics, and AI-based structural prediction, this study refines the stony coral biomineralization model from a simple list of “matrix components” to a coordinated network of export pathways, vesicle-mediated delivery, and ECM reworking. In doing so, it establishes *D. pertusum*, a reef-building, asymbiotic cold-water coral, as a novel symbiont-free model for dissecting the molecular mechanisms of coral calcification and provides a mechanistic framework for future functional assays and for understanding how this system may respond to ongoing ocean change.

### Limitations of the study

Limited sample availability of deep-sea corals reduces the replicability per analysis type. Destructive analysis limits cross-analysis sample availability.

## Resource Availability

### Lead contact

Dr. Fiorella Prada

### Materials availability

This study did not generate new unique reagents or deposited biological materials.

### Data and code availability

All data and code needed to evaluate and reproduce the conclusions in the paper are available in the paper, Supplementary Materials, and/or GitHub repository (); raw LC-MS/MS data are archived at MassIVE under with reviewer login = and password = ; these data will be publicly released upon publication of this manuscript.

## Acknowledgments

We thank the crew of the R/V Atlantis and the NOAA Ships Ronald H. Brown and Okeanos Explorer used to collect *D. pertusum* analyzed in this work. We thank Murray Roberts for contributing additional *D. pertusum* polyps for SEM, TGA, amino acid and density measurements. We thank the Rutgers Center for Advanced Biotechnology and Medicine Biological Mass Spectrometry Facility for generating and helping analyze LC–MS/MS data. We thank Ehud Zelzion from the Rutgers Office of Advanced Research Computing for providing guidance on bioinformatic analyses. *D. pertusum* sample collection was achieved through the Deep SEARCH project, funded by the Bureau of Ocean Energy Management (contract M17PC00009 to TDI Brooks International) and the NOAA Office of Ocean Exploration and Research (for ship time). Additional support came from the NOAA Deep-Sea Coral Research and Technology Program. This work was also supported by the Bennett L. Smith Endowment to PGF, funds from Rutgers University to FP and the Ministry of Innovation, Science & Technology, Israel to TM.

## Author contributions

Conceptualization: JD, TM, OL, FP; Resources: TM, EC, PGF, OL, FP; Methodology: JD, TM, LH, EC, SH, PGF, OL, FP; Investigation: JD, TM, SH, PGF, OL, FP; Formal Analysis: JD, TM, OL, FP; Validation: JD, TM, OL, FP; Data Curation: JD, FP; Visualization: JD, TM, OL, FP; Writing – original draft: JD, TM, LH, EC, SH, PGF, OL, FP; Writing: Review and Editing: JD, TM, LH, EC, SH, PGF, OL, FP; Supervision: EC, FP; Project Administration: JD, FP: Funding Acquisition: TM, EC, PGF, FP. All authors approve of the manuscript for review for publication.

## Supplemental information

**Document S1.** Supplemental text, figures, tables, and references.

**Data S1**. **Three hundred and eighty-two SOM proteins in *Desmophyllum pertusum* skeleton (.xlsx).** Proteins were detected by LC-MS/MS analysis, annotated in Blast2GO, and grouped according to their proposed function.

**Data S2. Thermogravimetric analysis of three coral skeletons (.xlsx).** *D. pertusum, S. pistillata,* and *O. patagonica* skeletons were heated from 35 to 600 ^°^C in N_2_. Weight loss is expressed as % values standardized to each sample’s starting mass. Raw data can be found at.

Data S3. *D. pertusum* reference protein database (.fasta). The full *D. pertusum* predicted proteome can be found at.

**Data S4 and S5. Non-redundant list of SOM proteins from *S. pistillata* and *O. patagonica* (.fasta).** Seventy-seven *S. pistillata* SOM proteins (*24, 27, 36*) and 75 *O. patagonica* SOM proteins (*47*) were determined by self-versus-self blast analysis.

**Data S6. Fifty-eight species included in OrthoFinder analysis.** 95.1% of genes across all taxa were assigned to 39,435 orthogroups.

**Data S7. Protein isoforms from 58 species assigned to orthogroup OG0001503 by OrthoFinder v3 (.fasta).** No isoforms from arthopods, choanoflagellates, chordates, echinoderms, fungi, nematodes, or poriferans were assigned to this orthogroup.

Data S8. *D. pertusum, S. pistillata,* and *O. patagonica* SOM proteins’ functional group and orthogroup assignments (.xlsx). All but 18 proteins were assigned to orthogroups by OrthoFinderv3.

## STAR Methods

### Experimental model details

#### Sample collection and preparation for protein extraction

Fragments of the cold-sea coral *Desmophyllum pertusum* (Linnaeus, 1758) were obtained as part of the Deep-Sea Exploration to Advance Research on Coral/Canyon/Cold-seep Habitats (Deep SEARCH) project. Corals were collected from two dives with the human-occupied vehicle (HOV) Alvin (AL4962 & AL4963) aboard the R/V Atlantis in August, 2018 (AT41), three dives with the remotely-operated vehicle (ROV) Jason-II (J2-1128, J2-1129, & J2-1138) aboard the NOAA Ship Ronald H. Brown in April, 2019 (RB1903), and two dives of the ROV Deep Discoverer (EX1806-7 & EX1903L2-10) on the NOAA Ship Okeanos Explorer in June, 2018 (EX1806) and June, 2019 (EX1903L2) ^46^. Individual samples of *D. pertusum* (Nordleksa reef, Norway; 150 m), *S. pistillata* (Eilat, Israel; 5 m), and *O. patagonica* (Sdot-Yam, Israel; 5 m) for (a) microscopy were stored in ethanol or for (b) organic matter content were oxidized and dried.

### Method details

#### Scanning Electron Microscopy

Individual coral samples were rinsed of ethanol then immersed in 1% sodium hypochlorite (NaClO) for 10 min to remove tissue, rinsed thoroughly with distilled water, and dried overnight. Fragments were vacuum coated with gold (for conductivity) prior to examination under a ZEISS SigmaTM Scanning Electron Microscope (Germany), by using an in-lens detector (2 kV, WD = 3–4 mm).

#### Thermogravimetric Analysis

Individual samples of *D. pertusum* (N = 1), *S. pistillata* (N = 3), and *O. patagonica* (N = 1) were subjected to heating in N_2_ at a rate of 20 ml/min and analyzed using Pyris software v12.1.1.0106 from 35 to 600 ^°^C in ∼0.17 ^°^C increments using PYRIS TGA (PerkinElmer, Shelton, CT, USA). Thermogravimetric analysis profiles were standardized to each sample’s starting mass, with weight loss expressed as % values (Figure S1; Data S2:). Sample sizes for TGA reflect the limited availability of cold-water coral material and of archived material for the comparative species. This sampling design is consistent with established practice in comparative biomineralization studies, where TGA characterization of coral skeletal organic matrices is typically reported on a single bulk sample per species (e.g.,^50,51^.

#### Amino Acid Analysis

Combined soluble and insoluble fractions from protein extractions of *D. pertusum* (N = 1), *S. pistillata* (N = 3), and *O. patagonica* (N = 1) skeletons were subjected to acid hydrolysis of peptide bonds at 100 °C in 6 N HCl. Samples were derivatized and immediately analyzed using GC-Trace 1310 MS-ISQLT equipped with IRMS-Delta V Advantage system (Thermo Scientific). Detected amino acids are presented as molar % (relative to all detected amino acids). By this method, Asn and Gln lose their terminal nitrogen during acid hydrolysis and are converted to Asp and Glu, respectively; they are therefore presented as Asx (Asp + Asn) and Glx (Glu + Gln)Glu, respectively ^134^. As for TGA, sample sizes for amino acid analysis follow established practice in comparative coral biomineralization studies, in which bulk amino acid composition is characterized on one specimen or a small pool per species (e.g.,^53^. Bulk amino acid composition and thermogravimetric profiles of coral SOM are species-typical and stable across specimens when samples are properly cleaned of endolithic and tissue-derived contamination (see Skeletal protein extraction and purification). Comparisons across species should nonetheless be regarded as preliminary, and future work expanding these analyses across additional specimens per species will be valuable.

#### Skeletal porosity, bulk density, and micro-density

Skeletal porosity, bulk density, and micro-density were quantified for *D. perusum* (N = 5), *O. patagonica* (N = 3), and *S. pistillata* (N = 2) using buoyant-weight (hydrostatic) measurements. After recording dry weight (DW), samples were vacuum-degassed in a drying chamber (3 h) to remove air from pores, then gradually saturated with MilliQ water and weighed in air to obtain saturated weight (SW). Buoyant weight (BW) was measured submerged using a hydrostatic balance (Ohaus Explorer Pro, ±0.0001 g) with a density determination kit.

Assuming water density ρ = 1 mg/mm³ (20 °C, 1 atm), we calculated:

Matrix volume: (V_MATRIX_ = (DW - BW)/ρ)
Pore volume: (V_PORES_ = (SW - DW)/ ρ)
Total (bulk) volume: (V_TOT_ = V_MATRIX_ + V_PORES_)

These volumes were used to derive ^135–137^:

Micro-density (mg/mm³): (DW / V_MATRIX_) (should not exceed 2.94 g cm⁻³, the density of pure aragonite;^138^
Bulk density (mg/mm⁻³): (DW / V_TOT_)
Porosity (%): ((V_PORES_ / V_TOT_) × 100), representing open (externally connected) pore space only.

#### Skeletal protein extraction and purification

A *Desmophyllum pertusum* fragment (∼2 g dry weight) was prepared for protein extraction following previous stony coral ^27^ and foraminifera (*135*) proteomics analyses. Briefly, fragments were oxidized with 20 mL 1:1 of 30% H_2_O_2_: 3% NaClO solution for 3 days, changing oxidizing solution once a day. Fragments were washed five times with ultra-pure water for one minute each time and dried at 50 °C overnight. Cleaned fragments were ground with a mortar and pestle to ≤ 150 μm diameter particles. Powder was oxidized in sterile Falcon brand tubes with 5 mL 1:1 of 30% H_2_O_2_: 3% NaClO while mixing at room temperature and incubated overnight in oxidizing solution. Powder was then washed in ultra-pure water three times to ensure that no organic residue remained on the mineral grains. In each cycle, the removal of the oxidizing or wash solution was performed by centrifugation at 5000 × g for 30 min at 25 °C. Cleaned powder was then dried overnight at 50 °C. We carried out all the described processes in a laminar flow biological hood (apart from oven drying) with all preparation tools and surfaces bleached to avoid contamination.

Cleaning was sufficient to remove contaminant proteins, as determined by sonicating the cleaned powder at room temperature in filter-sterilized phosphate buffered saline (PBS, pH 7.4) for 30 min, pelleting the powder at 5000 x g for 30 min at room temperature, concentrating the supernatant on 3 kDa Centricon YM-3 ultra-centrifugal filter units (Millipore) and running bichronoic acid assays (BCA) and 8–16% SDS-PAGE TGX Stain-Free gels (Bio-Rad) (Figure S3).

Approximately 1.5 g of cleaned skeletal powder was decalcified using a 16 cm-long osmotic tube for dialysis (MWCO = 3.5 kDa Thermo Scientific) with 5 mL of 0.2 µm filtered milli-Q water. The sealed membrane was placed into 2 L of 0.1 M CH_3_COOH (glacial acetic acid) solution with stirring for at least 72 h. The membrane containing the dissolved skeletal organic matrix (SOM) was dialyzed against 0.2 µm filtered milli-Q water until the final pH was about 6. The obtained aqueous solution containing the OM was centrifuged at 15,000 x g for 30 min at 4 °C to separate the soluble (SSOM) and the insoluble (ISOM) SOM fractions. SSOM and ISOM proteins were separated by SDS-PAGE and bands were visualized by silver staining (Pierce silver stain for mass spectrometry).

#### Liquid chromatography-tandem mass spectrometry (LC-MS/MS)

*Desmophyllum pertusum* samples were analyzed by LC-MS using Nano LC-MS/MS (Dionex Ultimate 3000 RLSCnano System, Thermofisher) interfaced with Eclipse (ThermoFisher) at the Rutgers Center for Advanced Biotechnology and Medicine (CABM). Three microliters of 12.5 µL in-gel digested using trypsin. Sample P was loaded on to a fused silica trap column (Acclaim PepMap 100, 75UMx2CM, ThermoFisher). After washing for 5 min at 5 µl/min with 0.1% TFA, the trap column was brought in-line with an analytical column (Nanoease MZ peptide BEH C18, 130A, 1.7 µm, 75umx250mm, Waters) for LC-MS/MS. Peptides were fractionated at 300 nL/min using a segmented linear gradient 4-15% B in 30 min (where A: 0.2% formic acid, and B: 0.16% formic acid, 80% acetonitrile), 15-25% B in 40min, 25-50% B in 44min, and 50-90% B in 11min. Solution B then returns at 4% for 5 minutes for the next run. The scan sequence began with an MS1 spectrum (Orbitrap analysis, resolution 120,000, scan range from M/Z 375–1500, automatic gain control (AGC) target 1E6, maximum injection time 100 ms). The top S (3 sec) duty cycle scheme were used to determine the number of MS/MS performed for each cycle. Parent ions of charge 2-7 were selected for MS/MS and dynamic exclusion of 60 seconds was used to avoid repeat sampling. Parent masses were isolated in the quadrupole with an isolation window of 1.2 m/z, automatic gain control (AGC) target 1E5, and fragmented with higher-energy collisional dissociation with a normalized collision energy of 30%. The fragments were scanned in Orbitrap with resolution of 15,000. The MS/MS scan ranges were determined by the charge state of the parent ion, but lower limit was set at 110 amu.

#### MS Database Search

The peak list of the LC-MS/MS were generated by Thermo Proteome Discoverer (v. 2.1) into MASCOT Generic Format (MGF) and searched against a common lab contaminants (CRAP) database using an in-house version of X!Tandem (GPM Fury) (Craig and Beavis 2004). Search parameters were the following: fragment mass error, 20 ppm; parent mass error, +/- 7 ppm; fixed modification, no fixed modification; variable modifications: methionine monooxidation for the primary search, asparagine deamination, tryptophan oxidation and dioxidation, methionine dioxidation, and glutamine to pyro-glutamine were considered at the refinement stage. Protease specificity: trypsin (C-terminal of R/K unless followed by P with 1 missed cleavage during the preliminary search and 5 missed cleavages during refinement. Minimum acceptable peptide and protein expectation scores were set at 10-2 and 10-4, respectively. The overall peptide false positive rate ^139^ was 0.07%.

#### Bioinformatic analysis

We used the *Desmophyllum pertusum* genome ^140^ as reference peptide databases (Data S3:) for the mass spectrometry analysis. Despite the inclusion of a contaminants database, several proteins likely of human origin were sequenced and attributed to *D. pertusum*. To filter out these potential contaminants from our final list of proteins, we BLASTed all sequences against the ‘Primates’ database in NCBI (taxa: 9443, Primates) using Blast2GO (number of blast hits = 20). We then examined NCBI-generated sequence alignments of *D. pertusum* versus Homo sapiens proteins with e-values lower than e^−50^. We removed from our final list of test proteins any sequences with: i) e-values lower than e^-100^, ii) percent similarity >80%, and three or more peptides each of seven or more amino acids in length that were identical between *D. pertusum* and humans. All remaining skeletal-specific proteins identified by the LC-MS/MS analysis were filtered to those with at least two peptides or at least one peptide with at least 10 spectra, and used for BLAST analysis against NCBI. Predicted proteins with similarities >40% and e-values <10^-10^ were retained. Transmembrane regions were predicted using the TMHMM server v2.0 ^141^. Signal peptides on the N-termini of proteins were predicted using SignalP 5.0 ^142,143^. Attachment of proteins to the exterior of the cell membrane by glycosylphosphatidylinositol (GPI) anchors was predicted using PredGPI ^144^. These computational analyses used program default settings and cut-offs.

Homology of *D. pertusum* SOM proteins with those of *S. pistillata* and *O. patagonica* was assessed via BLAST against their published skeletal proteomes. Additionally, a non-redundant list of published *S. pistillata* SOM proteins ^24,27,36^ was generated through self-versus-self BLAST to yield 77 proteins (Data S4), while the *O. patagonica* SOM proteome ^4^ yielded 75 proteins (Data S5) after BLASTing to the updated genome in NCBI ^59^. Gene ontology (GO) of all *D. pertusum, S. pistillata,* and *O. patagonica* SOM proteins was determined using BLAST2GO. Proposed protein function was then assigned based on GO terms and literature support for annotations.

We conducted an orthology analysis of known coral SOM proteins ^24,25,31,47^ within Metazoa - including 15 scleractinian proteomes - as well as fungi and choanoflagellates as outgroups. Most taxa were chosen to align with a previous orthology analysis ^47^; these were supplemented to ensure breadth and depth of both in- and out-groups yielding 58 genome-based proteomes (Data S6).

Reference genomes, annotated gene models, and/or select proteomes were downloaded from NCBI. For taxa without annotated gene model files in NCBI, genomes were annotated using BRAKER3 ^145^ in the European Galaxy server (Freiburg Galaxy Team; usegalaxy.edu) against taxonomically relevant proteomes. Annotated gene models were then reduced to single longest isoforms using the AGAT GTF/GFF analysis toolkit ^146^ in Galaxy. Reduced gene models were filtered and converted to predicted protein files using GFFRead ^147^ in Galaxy and then analyzed in OrthoFinder v3 using default settings. Quality of overall and species-specific orthology analysis was determined by percent of 1.3 million overall genes assigned to 39,435 orthogroups; this was found to be 95.1% overall and individual species gene assignment ranged from 74.3-99.4% (Data S6). The lower limit of %D+E in ’acidic’ proteins from coral skeleton was determined from proteins annotated as acidic SOMP or clustered with such proteins (Figure S4, Data S7). All scleractinian proteins falling in Orthogroup OG0001503, which contains CARP4/CARP5/SAARP1/SAARP2 and acidic SOMP, were clustered to 50% similarity in CD-Hit ^148^; the lowest %D+E for sequences in clusters that contained proteins annotated as acidic SOMP with a pI < 4.5 ^30^ was observed as 14.5% in *S. pistillata* (NCBI accession number XM_022926104.1). SOM proteins from *D. pertusum, S. pistillata,* and *O. patagonica* were assessed for patterns between functional group and orthogroup assignments (Figure 2, Data S8).

#### Fold Predictions and Comparisons

All identified SOM protein sequences were subjected to structural predictions via deep learning AI AlphaFold and ApfaFold3 ^124^. By using these predictions, AF3 has generated three-dimensional models which were further analyzed for their resemblance to other known folds available in the RCSB database using the DALI server ^149^. We further compared the predictions between the folds originated from *D. pertusum* and *S. pistillata* and found common folds as well as models that differ and are unique to each organism.

#### Key resources table

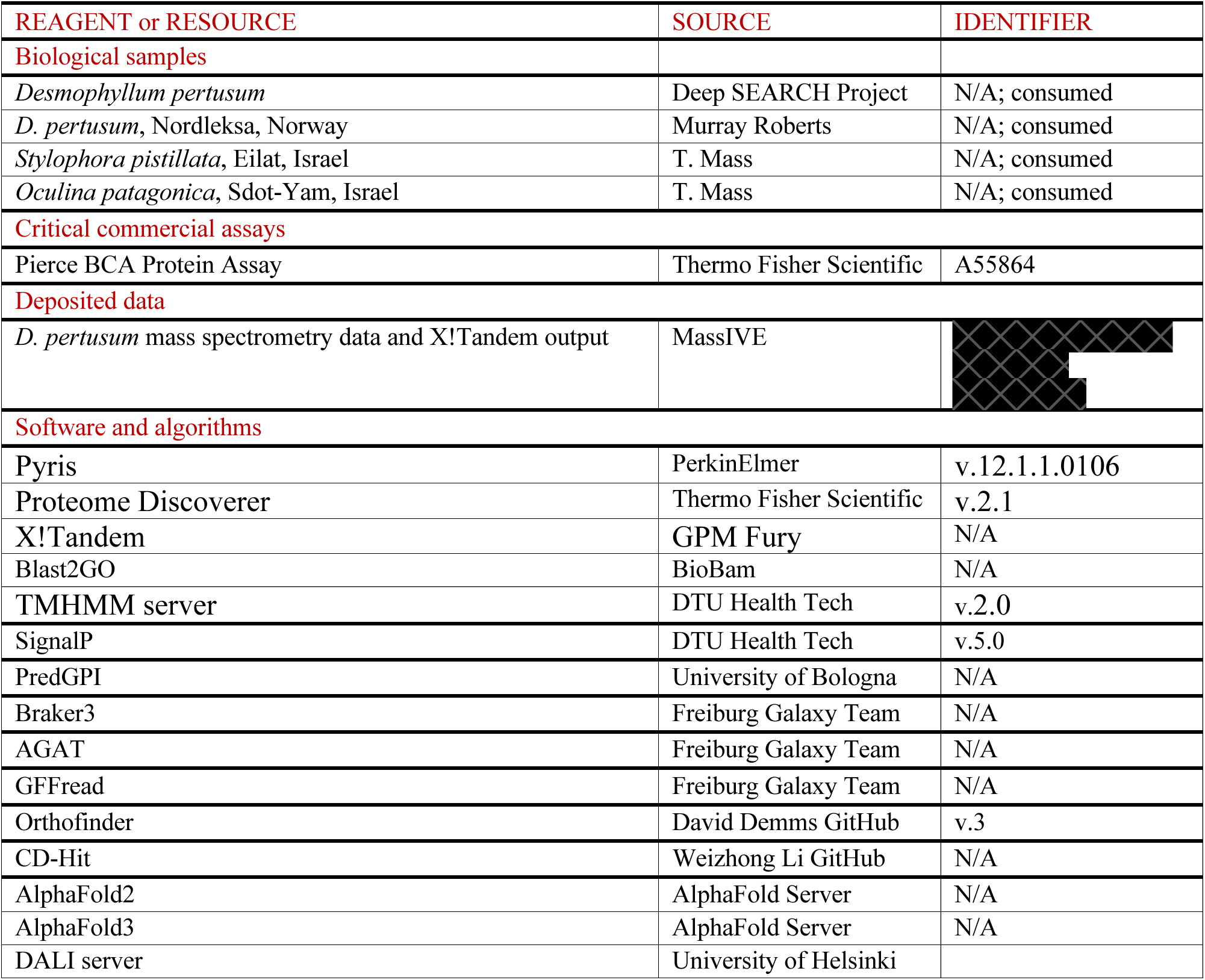

