## Supplementary Information for "Novel insights into conserved biomineralization mechanisms revealed from a cold-water scleractinian coral skeletal proteome"

**This PDF file includes:**

Supplementary Text
Figures S1 to S8
Table S1
Data S1 to S8
References (55, 150 to 155)

### Supplementary Text

Structural prediction description of species-specific SOM proteins

*Stylophora pistillata*

CARP1 (pI=4.12, D/E=36.9%) strongly resembles a calumenin with six putative $Ca^{2+}$ binding sites that are coordinated in an almost identical manner (Figure S5). Its D/E rich segment at the N-term region (residues 22-89; 68 of 347 19.6%) also could not be modeled by AF due to its exceptional sequence characteristics.

CARP2 (pI 4.76, 32.25% are D/E), CARP3 (pI=3.04, 50.3% D/E) and CARP6 (pI=3.5, D/E = 66%) contain very high DE composition, leaving them intrinsically disordered in the absence of cations by the AF3 analysis (*55*). Structures for these three acidic proteins are not shown, as intrinsically disordered regions are unresolved in AF3.

*Desmophyllum pertusum*

SAARP8 (pI=4.4 and D/E=16%), SAARP9 (pI=4.3 and D/E=17%), and SAARP10 (pI=3.7 and D/E=26%) contain very high DE composition, leaving them intrinsically disordered in the absence of cations by the AF3 analysis. As with CARP2, CARP3, and CARP6, we do not show these unresolved structures in AF3.

SAARP11 (pI=3.6, D/E=25%) exhibits an N terminus and a C terminus with very low pIDTT in AF3 prediction (Figure S6). The core of the protein folds similarly to α-galactosidase A with an $(\beta/\alpha)_8$ TIM barrel fold (48.3% sequence identity to the human protein). The enzyme α-GalA is regularly cleaving glycolipids. It is usually homodimeric and thus the prediction of SAARP11 was conducted on two polypeptide chains. However, there are indications that there are higher oligomerizations for α-Gal A in other species such as a tetramer in yeast [150] or *Mortierella vinacea* (fungi) [151] and a hexamer in *Thermus thermophilus* [152]. There are also alpha-Gal A in invertebrates such as mollusks, annelids, insects, and echinoderms but their oliogomerization state is not completely defined and can range between monomer and multimer.

SAARP13 (pI=2.7, D/E=40%) contains 729 residues and is highly acidic. AF3 prediction results in a unique core structure resembling antifreeze or pentapeptide-repeat proteins (Figure S7). The unique sequence contains a repeating DDSG motif that is accountable for the unique predicted topology. Other than the unpredicted parts (residues 1-109 and 553-729), it contains an extended highly acidic (more than 150 Å) cavity that can coordinate many $Ca^{2+}$ cations. It is plausible that the external surface, which is electronegative, will also bind available cations.

Another newly discovered protein unique to *D. pertusum* is annotated and shows a structural prediction as a protein disulfide isomerase (PDI; FUN_010535-T1). While PDIs generally consist of four domains (a-b-b'-a') - where domains 1 and 4 (a and a') are thioredoxin-like causing the CGHC active site motif to catalyze the formation, breakage, and rearrangement of disulfide bonds in nascent proteins within the endoplasmic reticulum - there is only one catalytic domain (Figure S8) in the *D. pertusum* SOM protein, which may allow this protein to retain some activity as has been observed in other similar proteins such as ERp4 with the *a-b-b'* [153] and ERp18 with the *a-b* [154,155].

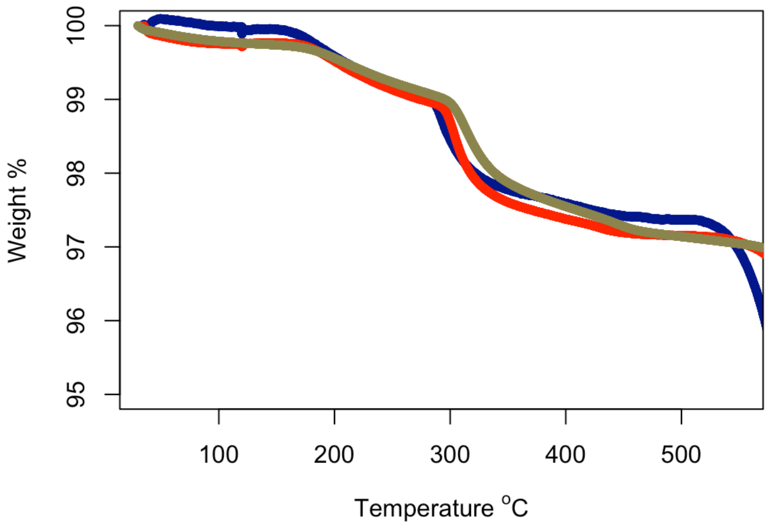


**Figure S1.** Thermogravimetric analysis of *D. pertusum* (blue), *S. pistillata* (red), and *O. patagonica* (dark khaki) skeletons. Weight change is presented as change from that at the initial temperature of 35 °C.

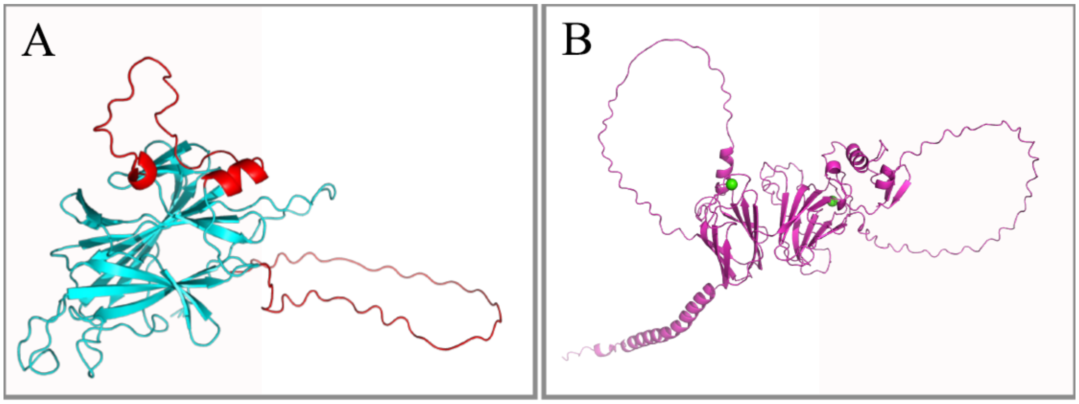


**Figure S2.** (A) CARP4/SAARP1 full prediction of fold with the D/E rich segments shown in red. The core of the β-sandwich is also typical to CARP5/SAARP2. (B) The prediction of full length SAARP12 from *D. pertusum* showing the two-β topologies in a tandem core with the undefined conformations of the D/E rich regions.

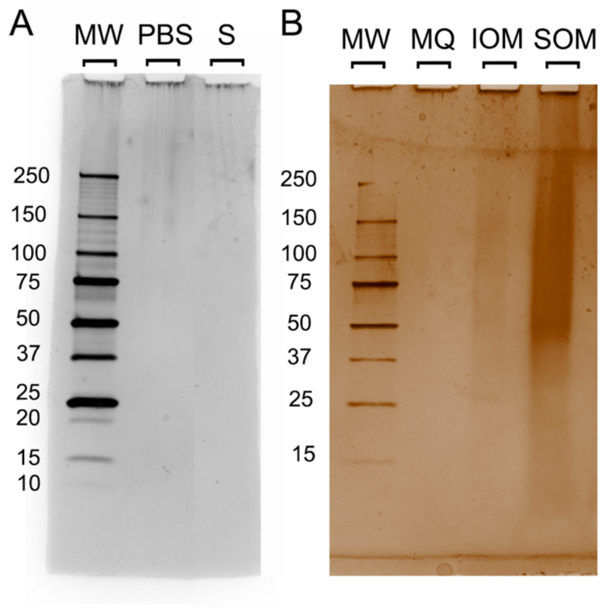


**Figure S3.** (A) Concentrated PBS that had been soaked on *Desmophyllum pertusum* powder oxidized three times and run on a 4 to 20% gel by SDS-PAGE. (B) *D. pertusum* organic matrix proteins separated by SDS/PAGE. Bands visualized by silver staining. MW (kDa): molecular weight, PBS: (phosphate buffered saline blank), S: (PBS soaked on cleaned *D. pertusum* powder and concentrated), MQ: (MilliQ ultrapure water), IOM: insoluble organic matrix, SOM: soluble organic matrix.

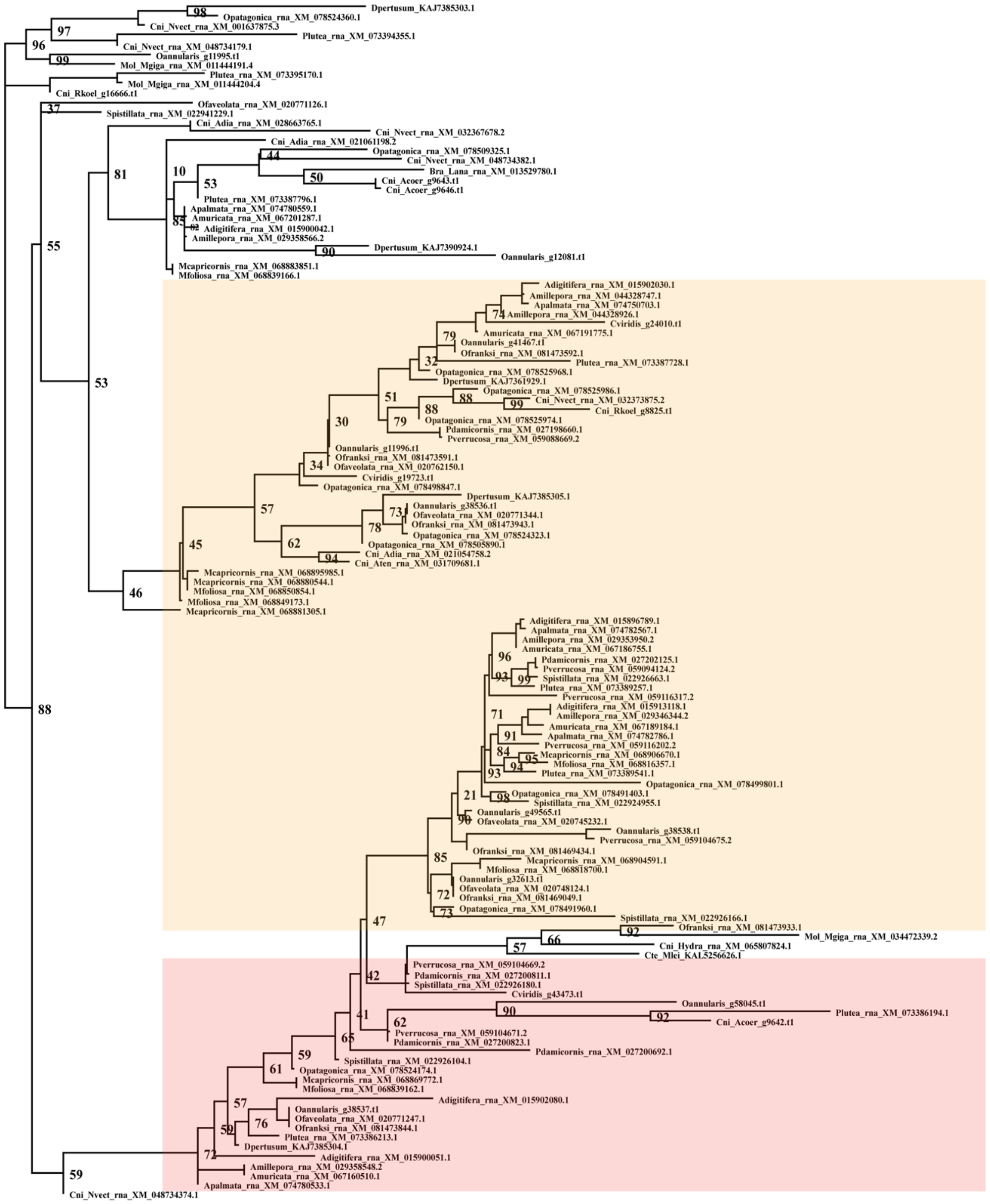


**Figure S4.** Orthogroup OG0001503 sequences as determined in OrthoFinder v3. This orthogroup encompasses all CARP4/SAARP1, CARP5/SAARP2, and acidic SOMP sequenced from stony coral skeleton previously and in the present study. Highlighted clusters comprise proteins identified as CARP4/SAARP1 and CARP5/SAARP2 (orange box) and acidic SOMP (pink box).

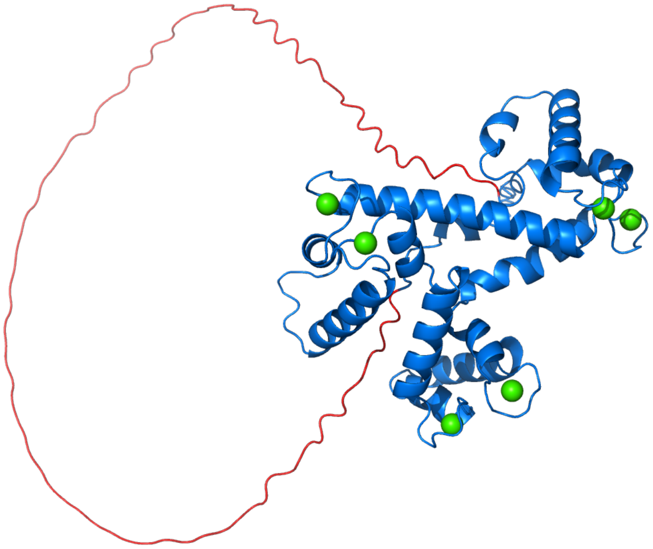

**Figure S5.** CARP1 prediction with the core fold similar to calumenin with six $Ca^{2+}$ binding sites (by similarity) shown in green. The N-term D/E rich region is shown in red and AF3 could not properly predict its conformation.

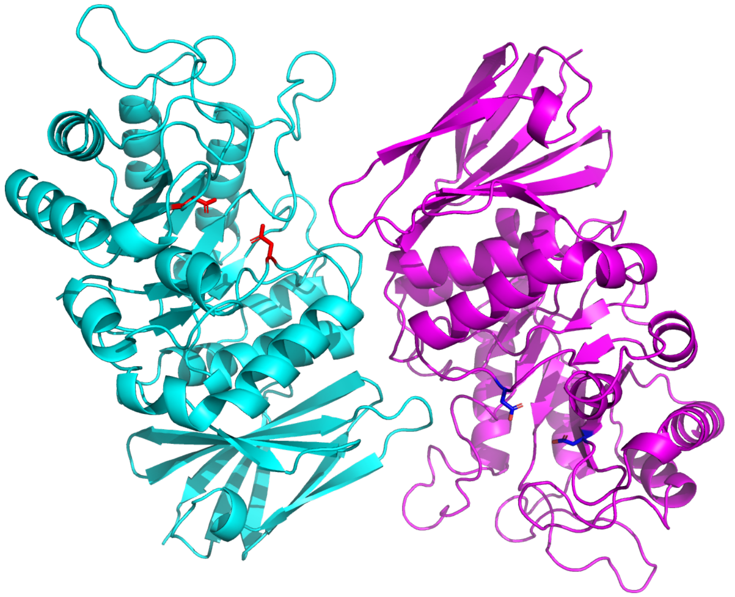

**Figure S6.** SAARP11 is an α-galactosidase A, shown here as a predicted dimer in magenta and cyan (without the disordered regions). This dimeric arrangement is tentative, and the oligomerization state in the coral may have a different multiplicity. Red and blue residues represent catalytic aspartic acids in the cyan and magenta monomers, respectively.

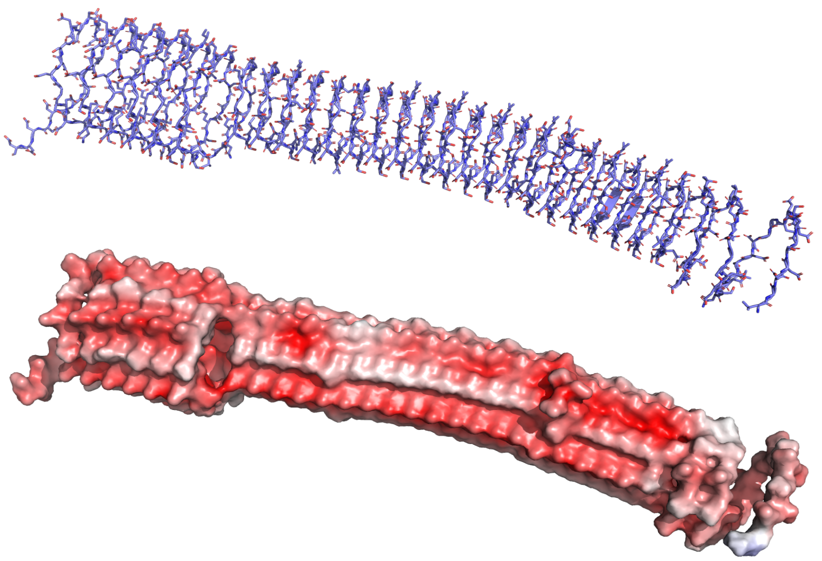

**Figure S7.** The core of SAAPR13 contains a unique fold 150 Å in length with many acidic residues pointing inwards into its core and outwards with the capability to bind numerous $Ca^{2+}$ cations (top). The bottom panel show a potential surface where red indicates electronegativity.

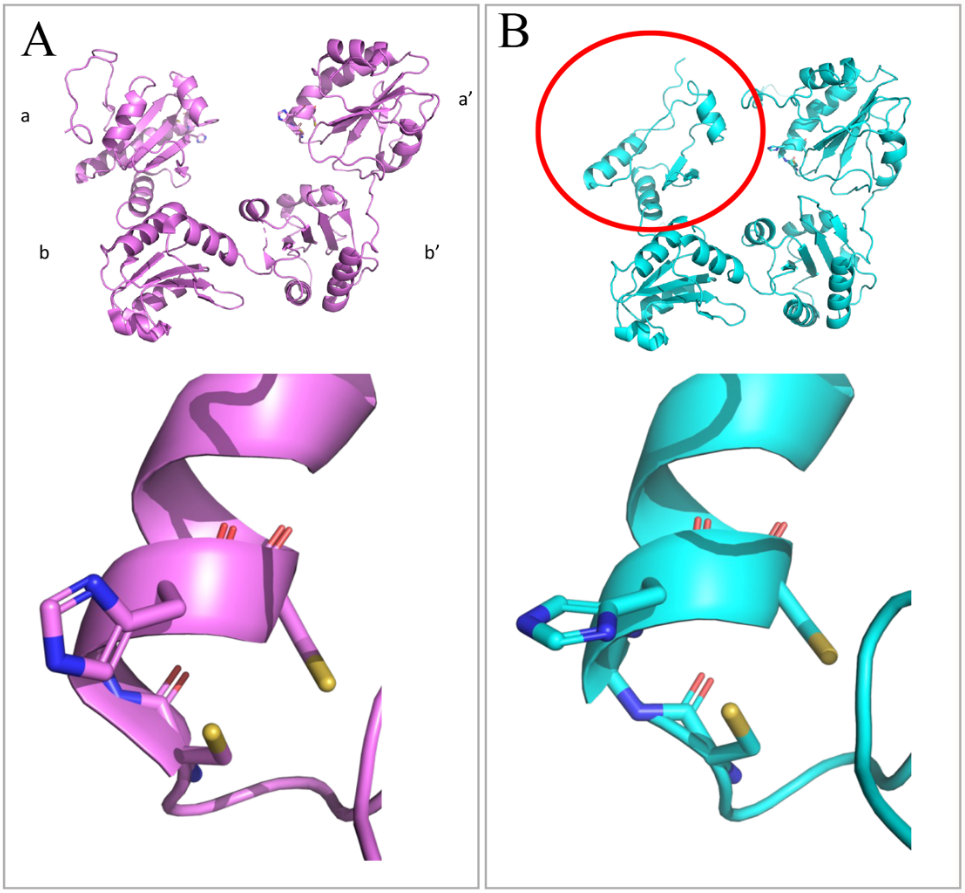


**Figure S8.** Protein disulfide-isomerase (PDI). (A) Structure of the human PDI (PDB entry 4EKZ, magenta) consisting of 4 domains (top panel). The *a* and *a'* contain the CGHC active site motifs (bottom panel). (B) Structure of *D. pertusum* PDI (FUN_010535-T1) without the D/E rich region, exhibiting 4 domains (cyan, top panel) but with only one CGHC active site in the fourth domain (bottom panel). The first domain (circled in red) does not contain the CGHC active site motif.

| | GC-MS | Genome | GC-MS | Genome | GC-MS | Genome |
|---|---|---|---|---|---|---|
| % | *D. pertusum* | | *S. pistillata* | | *O. patagonica* | |
| **ALA** | 14.97 | 7.48 | 7.81 | 6.80 | 12.03 | 7.46 |
| **GLY** | 24.77 | 8.16 | 9.33 | 9.47 | 25.84 | 9.39 |
| **VAL** | 9.79 | 8.60 | 15.83 | 8.01 | 9.78 | 7.81 |
| **LEU** | 4.90 | 9.13 | 8.92 | 9.47 | 5.97 | 8.92 |
| **ILE** | 4.63 | 6.54 | 9.48 | 6.31 | 5.00 | 6.22 |
| **THR** | 1.91 | 7.91 | 1.30 | 8.13 | 1.96 | 7.76 |
| **PRO** | 4.04 | 5.96 | 4.82 | 5.58 | 4.62 | 5.99 |
| **ASX** | 16.93 | 14.03 | 13.97 | 14.08 | 16.70 | 15.20 |
| **MET** | 0.08 | 2.31 | 0.37 | 2.06 | 0.12 | 2.18 |
| **GLX** | 12.61 | 11.72 | 13.04 | 12.38 | 12.94 | 11.69 |
| **PHE** | 2.67 | 4.69 | 5.23 | 4.98 | 2.99 | 4.65 |
| **LYS** | 1.38 | 8.06 | 7.01 | 7.52 | 1.00 | 6.79 |
| **TYR** | 0.94 | 4.01 | 1.37 | 3.76 | 0.80 | 4.04 |
| **TRP** | 0.38 | 1.41 | 1.52 | 1.46 | 0.25 | 1.90 |
| **Sum** | 100 | 100 | 100 | 100 | 100 | 100 |
| **Asx+Glx** | 29.54 | 25.74 | 27.01 | 26.46 | 29.64 | 26.89 |

**Table S1.** Relative amino acid composition (molar %) as determined by GC-MS analysis of hydrolyzed and derivatized SOM extracted from *D. pertusum, S. pistillata,* and *O. patagonica*, and compared with sequence-derived amino acid composition.

**Data S1**. **Three hundred and eighty-two SOM proteins in *Desmophyllum pertusum* skeleton (.xlsx).** Proteins were detected by LC-MS/MS analysis, annotated in Blast2GO, and grouped according to their proposed function.

**Data S2. Thermogravimetric analysis of three coral skeletons (.xlsx).** *D. pertusum, S. pistillata,* and *O. patagonica* skeletons were heated from 35 to 600 ˚C in $N_2$. Weight loss is expressed as % values standardized to each sample's starting mass. Raw data can be found at [redacted].

**Data S3.** ***D. pertusum*** **reference protein database (.fasta).** The full *D. pertusum* predicted proteome can be found at [redacted].

**Data S4 and S5. Non-redundant list of SOM proteins from** ***S. pistillata*** **and** ***O. patagonica*** **(.fasta).** Seventy-seven *S. pistillata* SOM proteins (*24, 27, 36*) and 75 *O. patagonica* SOM proteins (*47*) were determined by self-versus-self blast analysis.

**Data S6. Fifty-eight species included in OrthoFinder analysis.** 95.1% of genes across all taxa were assigned to 39,435 orthogroups.

**Data S7. Protein isoforms from 58 species assigned to orthogroup OG0001503 by OrthoFinder v3 (.fasta).** No isoforms from arthopods, choanoflagellates, chordates, echinoderms, fungi, nematodes, or poriferans were assigned to this orthogroup.

**Data S8.** ***D. pertusum, S. pistillata,*** **and** ***O. patagonica*** **SOM proteins' functional group and orthogroup assignments (.xlsx).** All but 18 proteins were assigned to orthogroups by OrthoFinder v3.
